# Targeting extracellular glycans: Tuning multimeric boronic acids for pathogen-selective killing of *Mycobacterium tuberculosis*

**DOI:** 10.1101/529743

**Authors:** Collette S. Guy, Matthew I. Gibson, Elizabeth Fullam

## Abstract

Innovative chemotherapeutic agents that are active against *Mycobacterium tuberculosis (Mtb*) are urgently required to control the tuberculosis (TB) epidemic. The *Mtb* cell envelope has distinct (lipo)polysaccharides and glycolipids that play a critical role in *Mtb* survival and pathogenesis and disruption of pathways involved in the assembly of the *Mtb* cell envelope are the primary target of anti-tubercular agents. Here we introduce a previously unexplored approach whereby chemical agents directly target the extracellular glycans within the unique *Mtb* cell envelope, rather than the intracellular biosynthetic machinery. We designed and synthesised multimeric boronic acids that are selectivity lethal to *Mtb* and function by targeting these structurally unique and essential *Mtb* cell envelope glycans. By tuning the number of, and distance between, boronic acid units high selectivity to *Mtb*, low cytotoxicity against mammalian cells and no observable resistance was achieved. This non-conventional approach may prevent the development of drug-resistance and will act as a platform for the design of improved, pathogen-specific, next generation antibiotics.

## Introduction

*Mycobacterium tuberculosis*, the causative agent of tuberculosis (TB), is the world’s leading cause of death from a single infectious agent claiming the lives of 1.7 million people annually.^1^ The incidence of drug resistant strains of *Mtb* are increasing at an alarming rate and include the emergence of *Mtb* that is not treatable with any of the current antibiotic regimens.^1^ Consequently, there is an urgent need for the development of innovative, next-generation anti-tubercular treatments which function by distinct mechanisms compared to the current drugs available. *Mtb* possesses a distinctive cell envelope that is uniquely complex and rich in a diverse range of unusual carbohydrates and lipids.^2^ The cell envelope has a fundamental role in the pathogenesis and virulence of *Mtb* and provides a highly efficient permeability barrier that prevents intracellular access to many antibiotics and severely complicates anti-tubercular treatment regimens. The core of the *Mtb* cell wall is comprised of three main components: a cross-linked peptidoglycan (PG) network, a highly branched arabinogalactan (AG) with both arabinose and galactose found in the furanose form, and long chain (C_60-90_) mycolic acids.^3^ The outer ‘myco-membrane’ contains a large array of distinct glycolipids and lipoglycans that are interspersed within this core and include phosphatidylinositol mannosides (PIMS), phthiocerol dimycocerosates (PDIMs), lipomannan (LM), lipoarabinomannan (LAM), mannose-capped LAM (ManLAM), sulfolipids and trehalose mono-and di-mycolates (TMM, TDM).^4^ The final component of the *Mtb* envelope is an outer capsule composed of polysaccharides, predominantly *α*-glucan, and proteins.^5^, ^6^ Intriguingly, many of these carbohydrates are specific to the *Mycobacterium* genus and are found in unusual conformations with distinct glycosidic linkages. Molecular pathways directly involved in the biosynthesis of the *Mtb* cell envelope have proven to be especially vulnerable to chemotherapeutic agents and include the front-line drugs isoniazid^7^ and ethambutol^8^ and second-line drugs ethionamide^7^ and D-cycloserine.^9, 10^ The current TB drug development portfolio capitalises on validated, druggable intracellular pathways involved in the synthesis of mycobacterial cell envelope components and include TBA-7371,^11^ BTZ043,^12^ PBTZ169^13^ and OPC-167832^14, 15^ that all kill *Mtb* by inhibition of the biosynthesis of arabinan.

The *Mtb* cell wall glycans are essential for its survival and pathogenesis ^4, 16^-^19^ and any disruption of the macromolecular complex can be lethal to the survival of the pathogen.^12, 19, 20^ Therefore, there is the tantalizing potential that these pathogen specific extracellular cell envelope glycans may, themselves, be viable therapeutic targets and afford a strategy for overcoming the intrinsic *Mtb* cell envelope barrier. To evaluate this idea further (Fig. 1), we selected to develop synthetic glycan receptors with selectivity for *Mtb* glycans as they are highly tunable and more stable under physiological conditions than biological counterparts.^21^ We selected to exploit the boronic acid pharmacophore that is an established glycan-binding functional group and forms bonds with cis 1,2 and 1,3 diols that are present in carbohydrates.^22^ In addition, a number of boron containing agents are compatible for use in humans with a number of FDA approved boron-drugs in clinical use in other disease areas ^23-25^.

**Fig. 1.**
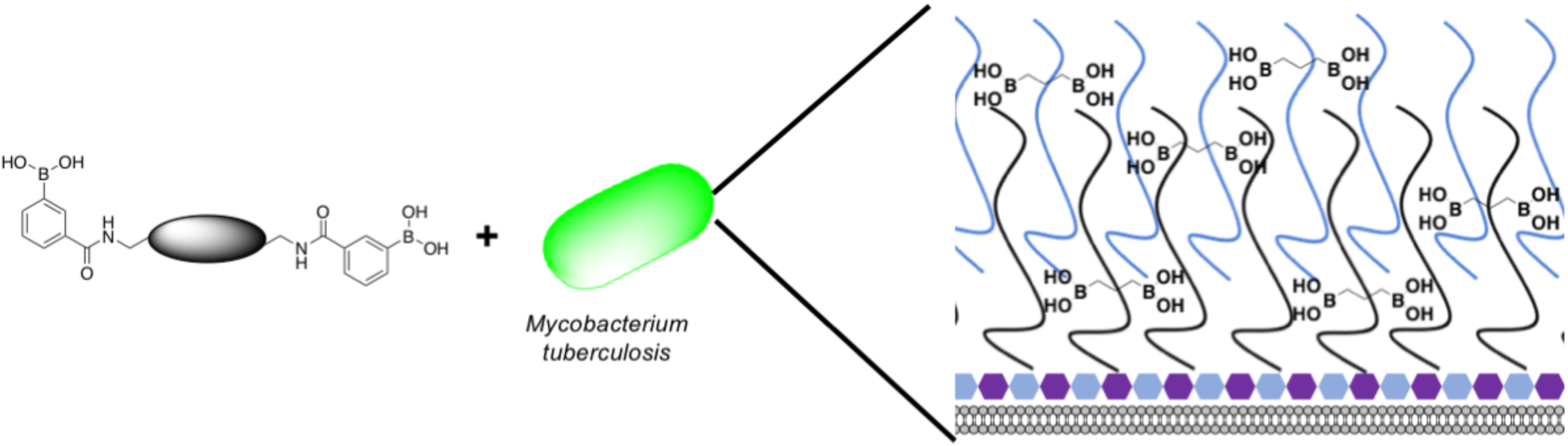
Overview of the approach used in this study. Illustration highlighting the design of multimeric boronic acids to specifically target the extracellular *Mtb* cell-envelope glycans. The complex *Mtb* cell envelope is simplified by black and blue lines, hexagons represent the peptidoglycan layer. This approach has several advantages over current strategies that include: no requirement for the molecule to cross the impenetrable *Mtb* cell wall barrier and the potential avoidance of drug efflux challenges.

We report here a new class of multimeric boronic acids which specifically kill *Mtb* through specific binding to *Mtb* cell envelope glycans. The most active compounds selectively kill mycobacteria over other strains of bacteria and exhibited low cytotoxicity to human cells. Whole-cell proteomics reveals a broad physiological stress response that does not result in the generation of resistance. The separation distance between the boronic acids was shown to be crucial for both activity and selectivity. These findings suggest that new classes of anti-tubercular therapies based on targeting the unique extracellular components are possible.

## Results and Discussion

### Design and synthesis of multimeric boronic acids

To explore the potential of multivalent boronic acid analogues to selectively target *Mtb* glycans, we designed and synthesised a panel of compounds (**1-8**) to evaluate the effect of the number and separation of boronic acids units to selectively target *Mtb* glycans and kill *Mtb*. A range of compounds bearing the glycan-targeting unit, 3-carboxy-phenyl boronic acid (3-CPBA), were synthesised (**4-8**) using acid chloride or carbodiimide coupling with the appropriate di-or tetra-PEG (polyethylene glycol) amine to provide flexibility of the spacer group between the relative position of the boronic acid functional groups (Supplementary schemes 1-3), giving the focused panel shown in Fig. 2 (details provided in Supplementary information, Schemes S1-S3, Figs S7-22). Systematic variation of the distance between the boronic acids (∼ 1.5 - 10 nm) was achieved using variable lengths of PEG (poly(ethylene glycol) diamines (**5-7**). Compound **4** had an ethyl linker as a further control, but longer alkyl chains were not soluble. For **8**, a first generation PAMAM (polyamidoamine) dendrimer core was synthesised and used to generate a tetrameric boronic acid (Supplementary information, Scheme S3). Systematic variation of the distance between the boronic acids (1.6 - 10 nm) was achieved using variable lengths of PEG (**5-7**). To ensure the boronic acids were accessible for glycan binding the Alizarin red (ARS) assay was employed to map the selectivity of the dimeric boronic acids against a panel of carbohydrates and were found to retain the same affinity trends as the monomeric boronic acids (Fig. S1).

**Fig. 2.**
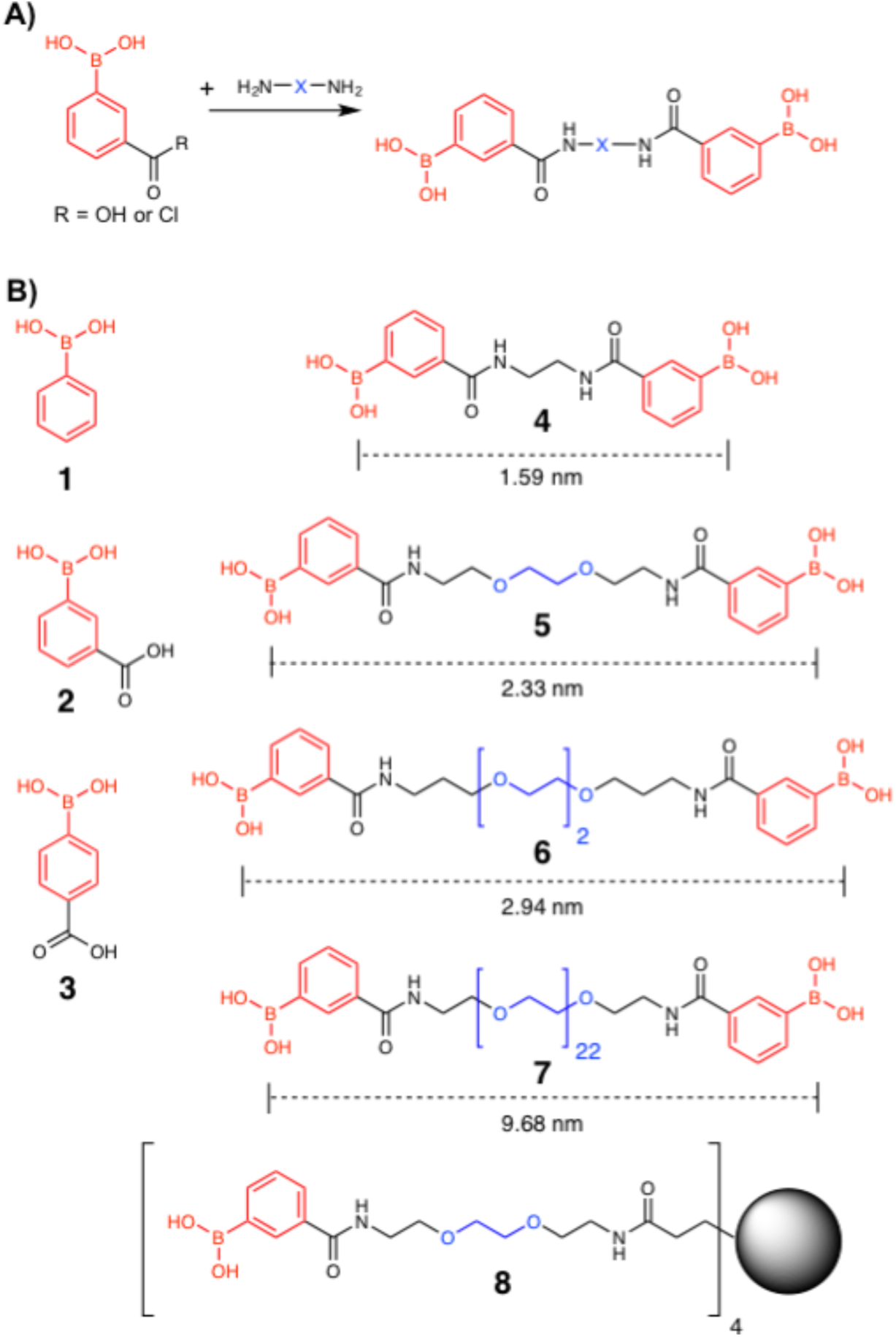
Boronic acid panel. A) Overview of synthetic approach; B) **1** – **3** were commercially available. **4** – **8** were synthesised as detailed in the Supplementary Information. Distances indicate the boron-boron distances assuming a fully extended chain: Angles of 109° and bond distances of 0.15 nm. Black circle is G1 PAMAM dendrimer.

### Determination of antibacterial potency

This library of glycan-targeting compounds was evaluated for antibacterial activity using the resazurin-reduction assay^26^ to determine the minimum inhibitory concentrations (MIC) (Fig. 3 and Table 1). The Gram-negative organisms *Escherichia coli* and *Pseudomonas putida* do not display the complex cell wall glycans of mycobacteria, and were tested alongside *Mycobacterium smegmatis, Mycobacterium bovis* BCG and also against *Mtb*. The monomeric boronic acids (**1-3**) displayed low antibacterial potency with no selective preference for Gram-negative or mycobacterial strains. Remarkably, exposure of the same strains to the dimeric boronic acids **4-6**, which vary in distance between the boronic acids from 12 – 23 atoms (1.6 – 3 nm), resulted in a dramatic decrease in the MICs (780 - 3100 αM against mycobacteria, Fig. 3, Table 1) and a corresponding increase in selectivity for mycobacteria compared to Gram-negative organisms. Notably, the dimeric compounds **4-6** were more effective against *Mtb* and *M. bovis* BCG than the widely used non-pathogenic model organism *M. smegmatis,* which is faster growing. Compound **7**, with a longer linker (∼ 10 nm) was ineffective, with higher MICs and loss of specificity for mycobacteria. Compound **8**, based on a 1^st^ generation poly(amidoamine) dendrimer, also showed mycobacterial selectivity with similar MIC values to the dimers **4-6 (**MIC 780 αM, against *Mtb*). These results indicate that an optimal spacer length of 1.6 – 3 nm (compounds **4-6**) between the two boronic acid moieties is optimal whereas longer lengths > 9 nm (**7**) reduce potency and fits a hypothesis of cell-wall glycan chelation.

**Table 1.** Bacterial susceptibility as measured by resazurin reduction microtitre plate assay

|  | MIC Mycobacteria (mM) |  |  | MIC Gram-negative (mM) |  |
| --- | --- | --- | --- | --- | --- |
|  | *Mycobacterium smegmatis* | *Mycobacterium bovis* BCG | *Mycobacterium tuberculosis* | *Escherichia coli* | *Pseudomonas putida* |
| **1** | 12.5-25 | 3.13 | 6.25-12.5 | 25 | 12.5-25 |
| **2** | 12.5-25 | 12.5 | 6.25-12.5 | >25 | 12.5-25 |
| **3** | 25 | 6.25 | 6.25 | >25 | 25 |
| **4** | 3.13 | 0.78 | 1.56-3.13 | >25 | >25 |
| **5** | 3.13 | 0.78-1.56 | 1.56-3.13 | >25 | >25 |
| **6** | 3.13 | 0.78-1.56 | 1.56-3.13 | >25 | >25 |
| **7** | 25 | >25 | - | >25 | >25 |
| **8** | 0.78 | 1.56 | 0.78-1.56 | >25 | >25 |

**Fig. 3.**
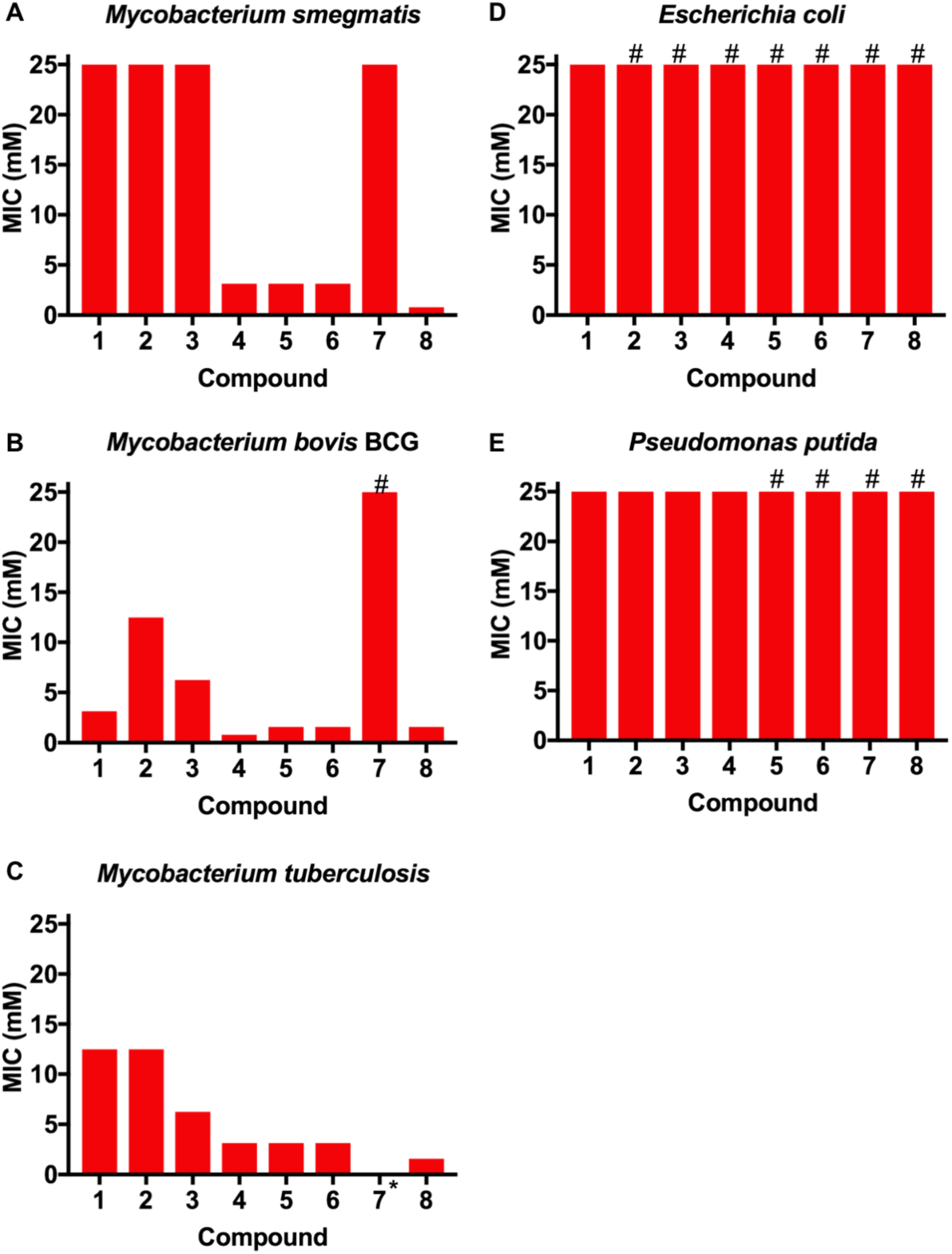
Antimicrobial activities of boronic acid derivatives **1-8** against A) *M. smegmatis,* B) *M. bovis* BCG, C) *M. tuberculosis*, D) *E. coli* E) *P. putida*. *, not tested; # represents MIC value greater than the maximum value tested (> 25 mM).

Minimal bactericidal concentrations (MBCs) were also determined against *Mtb* for **4-6**. The MBC data demonstrates that these boronic acid dimers are bactericidal against actively growing *Mtb* at concentrations of 6.25 mM (Table S1).

### *In vitro* cytotoxicity

The panel of boronic acids showed no significant cytotoxicity against human lung A549 cells with MIC_99_ values above 25 mM, and no lysis or agglutination of red blood cells was observed at concentrations as high as 25 mM (Table S2). The active dimers (**4-6**) and tetramer **8** therefore show higher selectivity for mycobacteria compared to the mammalian cells tested that have distinct cell-surface glycans.

### Lack of cross-resistance between dimeric boronic acid analogues and rifampicin and meropenem

To evaluate if the most active compounds (**4-6**) interacted with the front-line anti-tubercular agent rifampicin a checkerboard assay was used. No synergistic or antagonistic effects were noted with the sum of the fractional inhibitory index (ΣFIC) calculated as 2. Mono-boronic acids have been reported as having *β*-lactamase inhibitory activity ^27, 28^ and we therefore evaluated boronic acid dimers **4-6** for compound interactions with the *β*-lactam meropenem on *Mtb*. The ΣFIC for each combination was calculated ^29^ and found to be 1 for compound **4** and 0.6 for compounds **5-6** indicating no synergetic and importantly no antagonistic action on the growth inhibition, compared to the synergistic activity of meropenem in combination with the *β*-lactamase inhibitor sulbactam with a ΣFIC 0.3 (Table S3). These observations provide evidence that the multivalent boronic acids have a unique mechanism of action compared to the monomeric boronic acid *β*-lactamase transition state inhibitors and are not inhibitors of mycobacterial *β*-lactamase targets. Guided by the above results, we attempted to obtain resistant mutants of *M. bovis* BCG when plating on 5x MIC compound **5** but were unable to obtain mutants over a period of 3 months. This can be indicative that these dimeric compounds have a multifaceted mode of action. A low level of resistance has been found for antibiotics that target cell wall precursors including vancomycin, which took over 30 years for resistance to occur^30^, and the newly discovered texiobactin.^31^

### Identification of *Mtb* glycans as targets of dimeric boronic acids

To probe the selectivity and affinity of the 3-CPBA towards *Mtb* glycans biolayer interferometry (BLI) was employed (Fig. 4). A biotinylated boronic acid was synthesised (Scheme S4) and immobilised to streptavidin functionalised sensors. In control experiments against dextran and galactan that do not contain cis-diols, and are absent from the *Mtb* cell envelope, no binding was observed (Fig S4). A panel of isolated *Mtb* cell envelope components (Fig. 4) (additional information in supplementary information Figs S2-S4) were subsequently evaluated by BLI, revealing that the boronic acids interacted strongly with *Mtb* components that contain glycans with cis-diols (Fig. 4, Fig. S3): PG, AG, TMM, TDM, LAM and LM. *K*_*d*_ values were obtained using a steady-state model, giving values of 41 αg.mL^-1^ (PG), 4 αM (TDM) and 12 αM (TMM). Despite strong binding observed to AG we were unable to calculate the *K*_*d*_ value as saturation was not reached. In comparison, weak binding affinity towards PIM6 (which has six mannoses decorating the PIM unit) was observed, and very weak binding for PIM 1+2 (which have correspondingly lower degrees of glycosylation) and notably, no detectable binding for isolated mycolic acid methyl esters (MAMEs) and sulpholipid I which contains a sulphated trehalose moiety. This is consistent with a multivalent interaction between cell wall components that contain cis-diols and the boronic acid and clearly demonstrates that boronic acids have the necessary capacity to engage with these essential *Mtb* cell envelope constituents. As a final test, whole *E. coli* and (gamma-irradiated) *Mtb* cells were evaluated by BLI for binding to the boronic acid functionalised sensors. *Mtb* gave significantly faster rates and extent of binding compared to *E. coli (*Fig. 4C). Taken together, this BLI data supports a hypothesis that multimeric boronic acids selectively target mycobacterial cell wall glycans, leading to the observed bactericidal activity.

**Fig 4.**
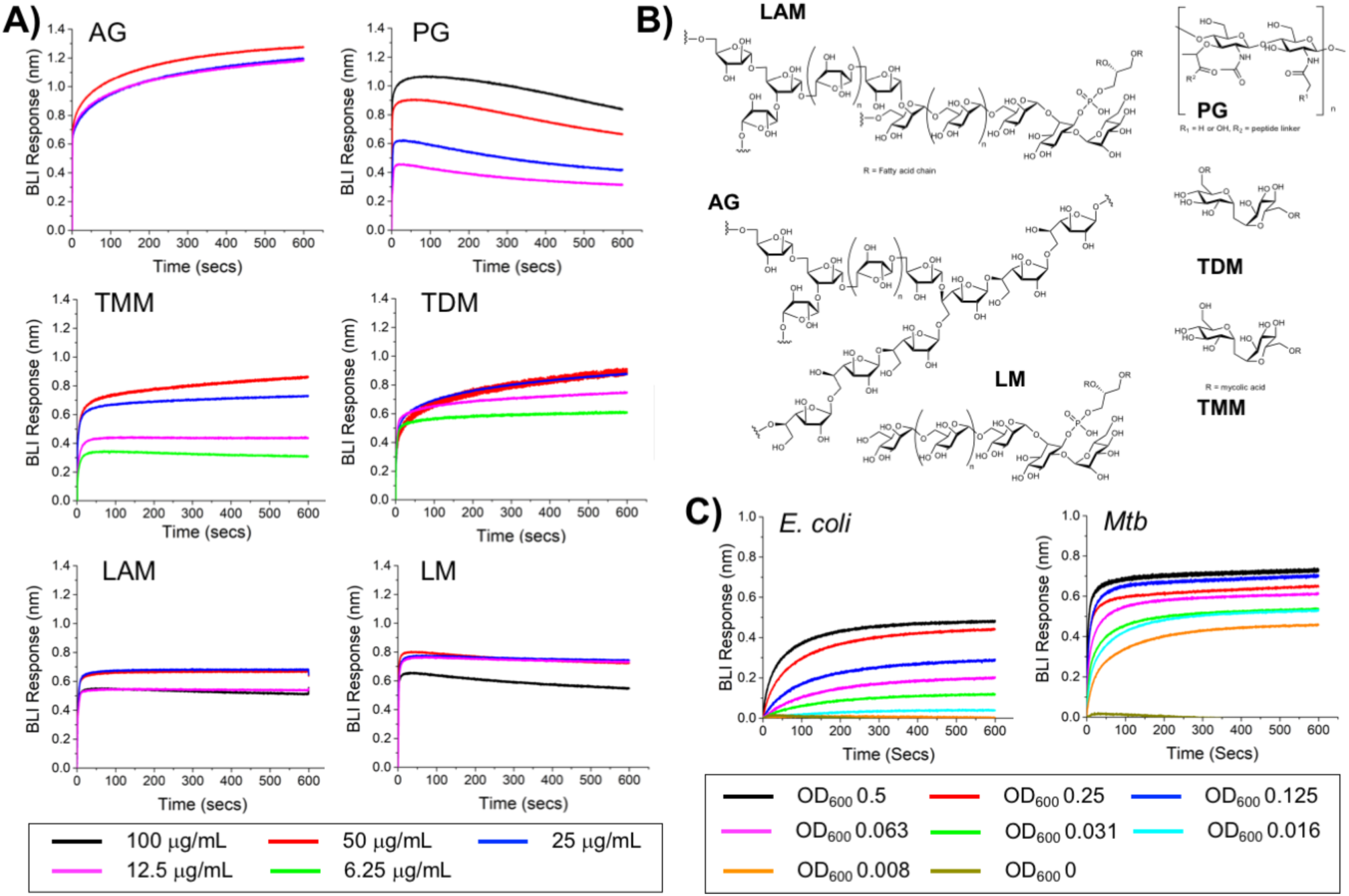
Biolayer interferometry analysis against 3-carboxy boronic acid functional sensor A) isolated *Mtb* cell envelope components, B) Structure of *Mtb* cell-envelope components, C) Whole *E. coli* and (gamma-irradiated) *Mtb* cells.

### Global protein expression response of *M. bovis* BCG to dimeric boronic acids

To gain physiological insight into the mode of action of these multimeric boronic acids, whole cell proteomics was employed. *M. bovis* BCG was exposed to 2x MIC of compound **6** and analysis of the whole cell protein expression profile at 3 hours, 24 hours and 48 hours was performed. Overall, we identified a total of >1,480 proteins and determined the differential expression after exposure to the boronic acid dimer (**6**) over time (Fig. S6). The proteins were sorted by their functional category based on the Tuberculist^32^ designations (Fig. 5, Fig. S6). A list for all the identified proteins, annotations and fold changes compared to controls at each time point are in Supplementary List S1.

**Fig. 5.**
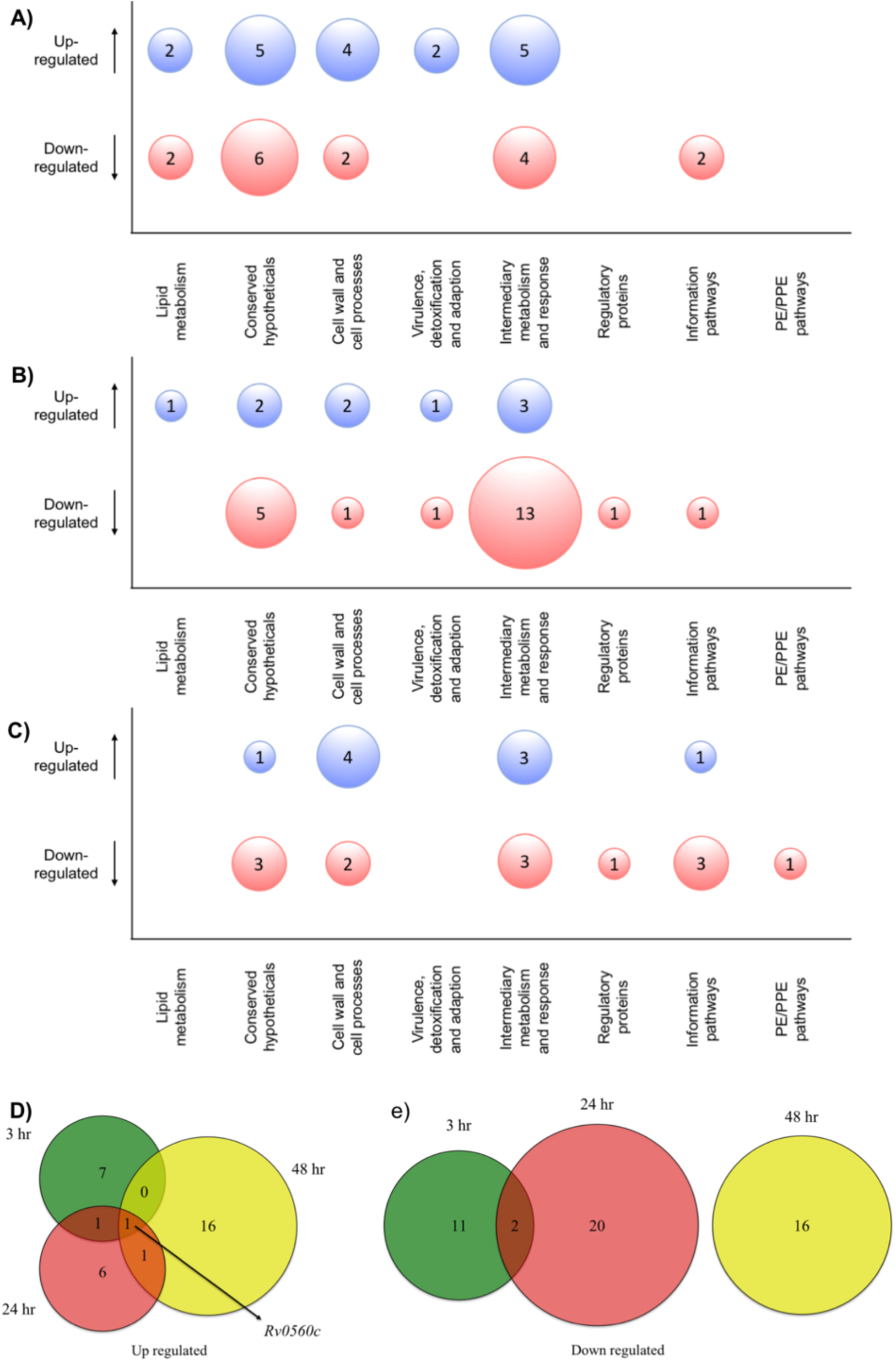
Whole cell proteomic analysis in response to **6**. Bubble plots show *Mtb* proteins that are up-or down-regulated at A) 3 hours, B) 24 hours and C) 48 hour. Venn diagrams indicate the number of proteins that are D) up-regulated or E) down-regulated after 3 hours (green), 24 hours (red) or 48 hours (yellow) exposure to **6**.

The proteins from two functional groups: category 3 (cell wall and cell processes) and category 7 (intermediary metabolism and respiration) were particularly affected, indicating that *Mtb* has a general stress response upon exposure of the dimeric boronic acids. In general, we found little overlap in the proteins that were either up-or down-regulated at the different time points (Fig. 5). However it was particularly notable that one identical protein that is involved in intermediary metabolism and respiration: Rv0560c, was upregulated >200 fold at 3 hours, 24 hours and 48 hours post-exposure to **6**. Rv0560c is a putative benzoquinone methyltransferase that has been associated with an increase in oxygen consumption by *Mtb* and is an indicator of a stress response.^32, 33^ Crucially, we detected no up-or down-regulation of penicillin binding proteins, *β*-lactamases, proteases and tRNA synthetase proteins (Supplementary List S1) that have been implicated as targets for mono-boronic acids ^27, 28, 34-36^ enabling us to rule out these intracellular targets, in-line with the meropenem checkerboard assays (Fig. S5, Table S3). Taken together this confirms that the multivalent presentation of boronic acids affords a new route to kill *Mtb*.

## Conclusions

In conclusion, here we have introduced a new approach to selectively kill *Mtb* by chelation of its unique cell wall glycans using multivalent boronic acids. This is conceptually distinct from existing drugs, which target defined intracellular pathways and hence must also permeate the *Mtb* cell envelope that confers intrinsic resistance to many antibiotics. The multivalent boronic acids were shown to selectively kill mycobacteria over other bacterial species. The distance between boronic acids was crucial with longer linkers reducing activity and selectivity. Two boronic acid units were optimal, with a tetramer (**8**) showing almost identical activity to the dimers. Biolayer interferometry revealed strong and selective interactions with isolated *Mtb* glycans and whole intact *Mtb* cells and whole cell proteomics identified a broad stress response rather than a single target, which may contribute to the lack of resistance observed. The multimeric display of boronic acids was crucial to their mechanism of action and distinct function compared to analogous monovalent boronic acids. This concept of inhibiting the extracellular glycans on *Mtb* presents a unique opportunity to develop pathogen specific agents and represents an important step in the identification of new TB drug targets.

### Conflicts of interest

The authors (CSG, MIG and EF) are named inventors on a patent application relating to this work.

## Acknowledgements

This work was supported by the Antimicrobial Resistance Cross Council Initiative supported by the seven Research Councils (grant no. MR/N006917/1), the EPSRC (grant no. EP/M027503/1) and the Royal Society (grant no. RG120405). EF holds a Sir Henry Dale Fellowship jointly funded by the Wellcome Trust and the Royal Society (grant no. 104193/Z/14/Z). The authors also acknowledge Trisha Bailey and Juan Hernandez-Fernaud for technical assistance and the WPH Proteomics Facility RTP, University of Warwick, and for assistance with the proteomics studies. *Mycobacterium tuberculosis* H37Rv cell envelope components and gamma irradiated whole cells were obtained through BEI Resources, NIAID, NIH.

## Notes and references

1. WHO Global Tuberculosis Report, http://www.who.int/tb/publications/global_report/en/).

2. P. J. Brennan and H. Nikaido, Annu Rev Biochem, 1995, 64, 29–63.

3. P. J. Brennan, Tuberculosis (Edinb), 2003, 83, 91–97.

4. M. Jackson, Cold Spring Harb Perspect Med, 2014, 4.

5. A. Lemassu and M. Daffe, Biochem. J., 1994, 297 (Pt 2), 351–357.

6. A. Ortalo-Magne, M. A. Dupont, A. Lemassu, A. B. Andersen, P. Gounon and M. Daffe, Microbiology, 1995, 141 (Pt 7), 1609–1620.

7. A. Banerjee, E. Dubnau, A. Quemard, V. Balasubramanian, K. S. Um, T. Wilson, D. Collins, G. de Lisle and W. R. Jacobs, Jr., Science, 1994, 263, 227–230.

8. K. Mikusova, R. A. Slayden, G. S. Besra and P. J. Brennan, Antimicrob Agents Chemother, 1995, 39, 2484–2489.

9. J. L. Strominger, E. Ito and R. H. Threnn, J Am Chem Soc, 1960, 82, 998–999.

10. G. A. Prosser and L. P. de Carvalho, ACS Med Chem Lett, 2013, 4, 1233–1237.

11. M. Chatterji, R. Shandil, M. R. Manjunatha, S. Solapure, V. Ramachandran, N. Kumar, R. Saralaya, V. Panduga, J. Reddy, K. R. Prabhakar, S. Sharma, C. Sadler, C. B. Cooper, K. Mdluli, P. S. Iyer, S. Narayanan and P. S. Shirude, Antimicrob Agents Chemother, 2014, 58, 5325–5331.

12. V. Makarov, G. Manina, K. Mikusova, U. Mollmann, O. Ryabova, B. Saint-Joanis, N. Dhar, M. R. Pasca, S. Buroni, A. P. Lucarelli, A. Milano, E. De Rossi, M. Belanova, A. Bobovska, P. Dianiskova, J. Kordulakova, C. Sala, E. Fullam, P. Schneider, J. D. McKinney, P. Brodin, T. Christophe, S. Waddell, P. Butcher, J. Albrethsen, I. Rosenkrands, R. Brosch, V. Nandi, S. Bharath, S. Gaonkar, R. K. Shandil, V. Balasubramanian, T. Balganesh, S. Tyagi, J. Grosset, G. Riccardi and S. T. Cole, Science, 2009, 324, 801–804.

13. V. Makarov, B. Lechartier, M. Zhang, J. Neres, A. M. van der Sar, S. A. Raadsen, R. C. Hartkoorn, O. B. Ryabova, A. Vocat, L. A. Decosterd, N. Widmer, T. Buclin, W. Bitter, K. Andries, F. Pojer, P. J. Dyson and S. T. Cole, EMBO Mol Med, 2014, 6, 372–383.

14. M. Matsumoto, H. Hashizume, T. Tomishige, M. Kawasaki, H. Tsubouchi, H. Sasaki, Y. Shimokawa and M. Komatsu, PLoS Med, 2006, 3, e466.

15. K. Mdluli, T. Kaneko and A. Upton, Cold Spring Harb Perspect Med, 2015, 5.

16. M. Daffe and P. Draper, Adv Microb Physiol, 1998, 39, 131–203.

17. O. Neyrolles and C. Guilhot, Tuberculosis (Edinb), 2011, 91, 187–195.

18. S. Pitarque, G. Larrouy-Maumus, B. Payre, M. Jackson, G. Puzo and J. Nigou, Tuberculosis (Edinb), 2008, 88, 560–565.

19. K. A. Abrahams and G. S. Besra, Parasitology, 2018, 145, 116–133.

20. L. Favrot and D. R. Ronning, Expert Rev Anti Infect Ther, 2012, 10, 1023–1036.

21. J. Arnaud, A. Audfray and A. Imberty, Chem Soc Rev, 2013, 42, 4798–4813.

22. H. G. Kuivila, A. H. Keough and E. J. Soboczenski, J. Org. Chem., 1954, 19, 780–783.

23. P. Hunter, EMBO Rep, 2009, 10, 125–128.

24. J. Adams and M. Kauffman, Cancer Invest, 2004, 22, 304–311.

25. B. E. Elewski, R. Aly, S. L. Baldwin, R. F. Gonzalez Soto, P. Rich, M. Weisfeld, H. Wiltz, L. T. Zane and R. Pollak, J Am Acad Dermatol, 2015, 73, 62–69.

26. J. C. Palomino, A. Martin, M. Camacho, H. Guerra, J. Swings and F. Portaels, Antimicrob Agents Chemother, 2002, 46, 2720–2722.

27. J. Brem, R. Cain, S. Cahill, M. A. McDonough, I. J. Clifton, J. C. Jimenez-Castellanos, M. B. Avison, J. Spencer, C. W. Fishwick and C. J. Schofield, Nat Commun, 2016, 7, 12406.

28. S. G. Kurz, S. Hazra, C. R. Bethel, C. Romagnoli, E. Caselli, F. Prati, J. S. Blanchard and R. A. Bonomo, ACS Infect Dis, 2015, 1, 234–242.

29. B. Lechartier, R. C. Hartkoorn and S. T. Cole, Antimicrob Agents and Chemother, 2012, 56, 5790–5793.

30. R. Leclercq, E. Derlot, J. Duval and P. Courvalin, N Engl J Med, 1988, 319, 157–161.

31. L. L. Ling, T. Schneider, A. J. Peoples, A. L. Spoering, I. Engels, B. P. Conlon, A. Mueller, T. F. Schäberle, D. E. Hughes, S. Epstein, M. Jones, L. Lazarides, V. A. Steadman, D. R. Cohen, C. R. Felix, K. A. Fetterman, W. P. Millett, A. G. Nitti, A. M. Zullo, C. Chen and K. Lewis, Nature, 2015, 517, 455.

32. A. Kapopoulou, J. M. Lew and S. T. Cole, Tuberculosis (Edinb), 2011, 91, 8–13.

33. Z. Sun, S. J. Cheng, H. Zhang and Y. Zhang, FEMS Microbiol Lett, 2001, 203, 211–216.

34. S. T. Cahill, R. Cain, D. Y. Wang, C. T. Lohans, D. W. Wareham, H. P. Oswin, J. Mohammed, J. Spencer, C. W. Fishwick, M. A. McDonough, C. J. Schofield and J. Brem, Antimicrob Agents Chemother, 2017, 61.

35. W. Moreira, G. J. Ngan, J. L. Low, A. Poulsen, B. C. Chia, M. J. Ang, A. Yap, J. Fulwood, U. Lakshmanan, J. Lim, A. Y. Khoo, H. Flotow, J. Hill, R. M. Raju, E. J. Rubin and T. Dick, MBio, 2015, 6, e00253–00215.

36. F. L. Rock, W. Mao, A. Yaremchuk, M. Tukalo, T. Crepin, H. Zhou, Y. K. Zhang, V. Hernandez, T. Akama, S. J. Baker, J. J. Plattner, L. Shapiro, S. A. Martinis, S. J. Benkovic, S. Cusack and M. R. Alley, Science, 2007, 316, 1759–1761.

