## Supplementary information for "Targeting extracellular glycans: Tuning multimeric boronic acids for pathogen-selective killing of *Mycobacterium tuberculosis*"

### Table of Contents

**Supplementary Results** 4

Scheme S1. Reaction scheme for the synthesis of boronic acids dimers **4** – **6** for 3-carboxyphenylboronic acid. 4

Scheme S2. Reaction scheme for the synthesis of boronic acid dimer **7** from 3-carboxyphenylboronic acid. 5

Scheme S3. Reaction scheme for the synthesis of 2,2'-(ethylenedioxy)bis(ethylamine) G1 PAMAM tetra(3-phenylboronic acid)amide dendrimer **8** 6

Scheme S4. Reaction scheme for the synthesis of biotinamido(hexanamido)(1,3-phenylboronic acid) **11**. 7

Figure S1. Alizarin red assay. 8

Figure S2. Structures of *Mtb* cell envelope components. 9

Figure S3. Biolayer interferometry response to functionalised boronic acid sensors. 10

Figure S4. Biolayer interferometry response to functionalised boronic acid sensors to glycans not found in the mycobacterial cell wall. 11

Figure S5. Representative images of checkerboard assays. 12

Figure S6. Functional distribution of total proteins identified by LC-MS in response to compound **6**. 13

Table S1. Minimum bactericidal concentrations of the dimeric boronic acids **4 - 6** against *Mycobacterium tuberculosis* H37Rv. 14

Table S2. Cytotoxicity testing of compounds **1-8** against human lung epithelium cell line and ovine blood 15

Table S3. MIC values against *M. tuberculosis* H37Rv and corresponding interaction profiles with rifampicin and meropenem as determined by REMA checkerboard assays. 16

**Supplementary Chemical and Synthetic protocols** 17

General Information and Procedures 17

Synthetic procedures 19

*N,N'*-(ethylenediamine)bis(3-phenylboronic acid)amide (**4**) 19

*N,N'*-(2,2-(ethylenedioxy)bis(ethylamine))bis(3-phenylboronic acid)amide (**5**) 19

*N,N'*-(4,7,10-trioxa-1,13-tridecanediamine)bis(3-phenylboronic acid)amide (**6**) 20

*N,N'*-(polyoxyethylene bis(amine))bis(3-phenylboronic acid)amide (**7**) 20

2,2'-(Ethylenedioxy)bis(ethylamine) G0.5 PAMAM dendrimer (**9**) 21

2,2'-(Ethylenedioxy)bis(ethylamine) G1 PAMAM dendrimer (**10**) 21

2,2'-(Ethylenedioxy)bis(ethylamine) G1 PAMAM tetra(3-phenylboronic acid)amide dendrimer (**8**) 22

Biotinamido(hexanamido)(1,3-phenylboronic acid) (**11**) 23

**Supplementary Biological protocols** 24

Bacterial strains, cell lines, culture conditions and chemicals 24

*Mycobacterium tuberculosis* H37Rv reagents 24

Alizarin Red S assay 24

Determination of minimum inhibitory concentrations 25

Determination of the minimum bactericidal (MBC) activity 25

Cytotoxicity assay 25

Haemolysis assay 25

Agglutination assay 26
Determination of compound interactions using a REMA checkerboard assay 26
Resistance studies 27
Biolayer Interferometry 27
Preparation of samples for proteomics 28
NanoLC-ESI-MS/MS Analysis 28
Data Analysis 28
Data processing and annotation 29
**References** 30
**NMR spectra** 31
Figure S7. $^{1}$H NMR spectrum of compound **4** 31
Figure S8. $^{13}$C DEPT NMR spectrum of compound **4** 32
Figure S9. $^{1}$H NMR spectrum of compound **5** 33
Figure S10. $^{13}$C DEPT NMR spectrum of compound **5** 34
Figure S11. $^{1}$H NMR spectrum of compound **6** 35
Figure S12. $^{13}$C DEPT NMR spectrum of compound **6** 36
Figure S13. $^{1}$H NMR spectrum of compound **7** 37
Figure S14. $^{13}$C DEPT NMR spectrum of compound **7** 38
Figure S15. $^{1}$H NMR spectrum of compound **9** 39
Figure S16. $^{13}$C DEPT NMR spectrum of compound **9** 40
Figure S17. $^{1}$H NMR spectrum of compound **10** 41
Figure S18. $^{13}$C DEPT NMR spectrum of compound **10** 42
Figure S19. $^{1}$H NMR spectrum of compound **8** 43
Figure S20. $^{13}$C DEPT NMR spectrum of compound **8** 44
Figure S21. $^{1}$H NMR spectrum of compound **11** 45
Figure S22. $^{13}$C DEPT NMR spectrum of compound **11** 46

#### Supplementary Results

**Scheme S1**. Reaction scheme for the synthesis of boronic acid dimers **4 - 6** from 3-carboxyphenylboronic acid.

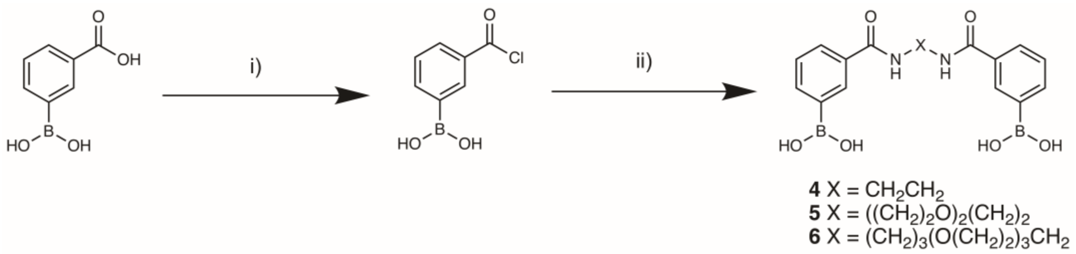


*Reagents and conditions:* i) 2.4 eq. oxalyl chloride, 0.2 eq. DMF, THF, 0 °C – room temperature, 2h. ii) 0.45 eq. diamine, 1 eq. pyridine, DMF, 0 °C – room temperature, 16h.

**Scheme S2**. Reaction scheme for the synthesis of boronic acid dimer **7** from 3-carboxyphenylboronic acid.

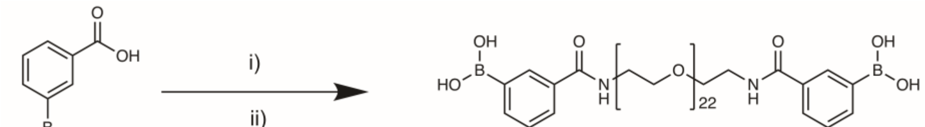


**7**

*Reagents and conditions:* i) 1.5 eq. EDCI, 0.5 eq. DMAP, DMF, 60 °C, 30 mins. ii) 0.45 eq. polyoxyethylene bis(amine) MW 1000 $g.mol^{-1}$, 60 °C, 16h.

**Scheme S3.** Reaction scheme for the synthesis of 2,2'-(ethylenedioxy)bis(ethylamine) G1 PAMAM tetra(3-phenylboronic acid)amide dendrimer **8**.

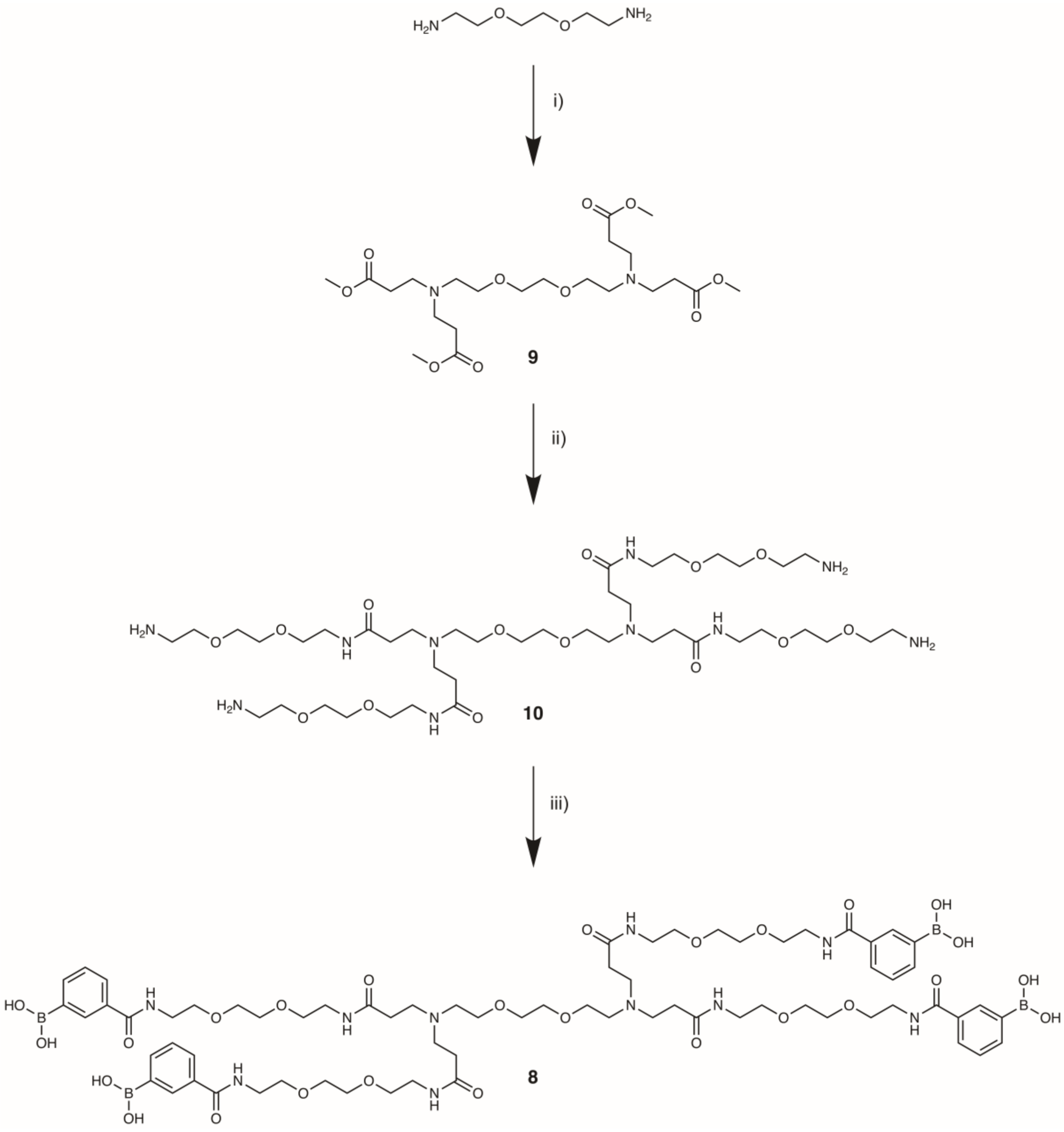


*Reagents and conditions:* i) 5 eq. methyl acrylate, methanol, 0 °C – room temperature, 96h. ii) 80 eq. 2,2'-(ethylenedioxy)bis(ethylamine), methanol, -30 °C – room temperature, 9 days. iii) 25 eq. (3-(chlorocarbonyl)phenyl)boronic acid, 25 eq. pyridine, DMAC, 0 °C – room temperature, 48h.

**Scheme S4.** Reaction scheme for the synthesis of biotinamido(hexanamido)(1,3-phenylboronic acid) (**11**).

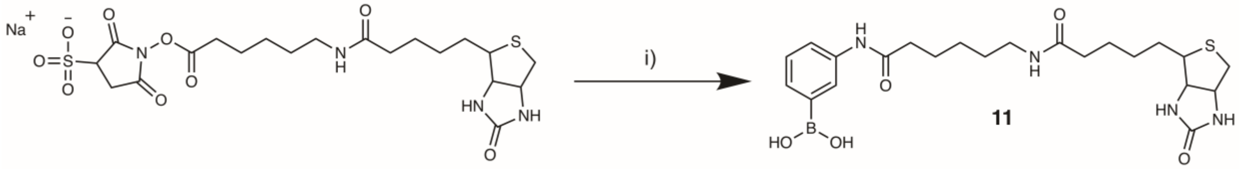


*Reagents and conditions:* i) 1 eq. 3-aminophenylboronic acid hydrochloride, 1.05 eq. triethylamine, DMF/water 1:4, room temperature, 16h.

**Figure S1.** Alizarin red assay. (a) 3-carboxyphenyl boronic acid **2**, (b) *N*,*N'*-(ethylenediamine)bis(3-phenylboronic acid)amide **4**, (c) *N*,*N'*-(2,2-(ethylenedioxy)bis-(ethylamine))bis(3-phenylboronic acid)amide **5**, (d) *N*,*N'*-(4,7,10-trioxa-1,13-tridecane-diamine)bis(3-phenylboronic acid)amide **6**.

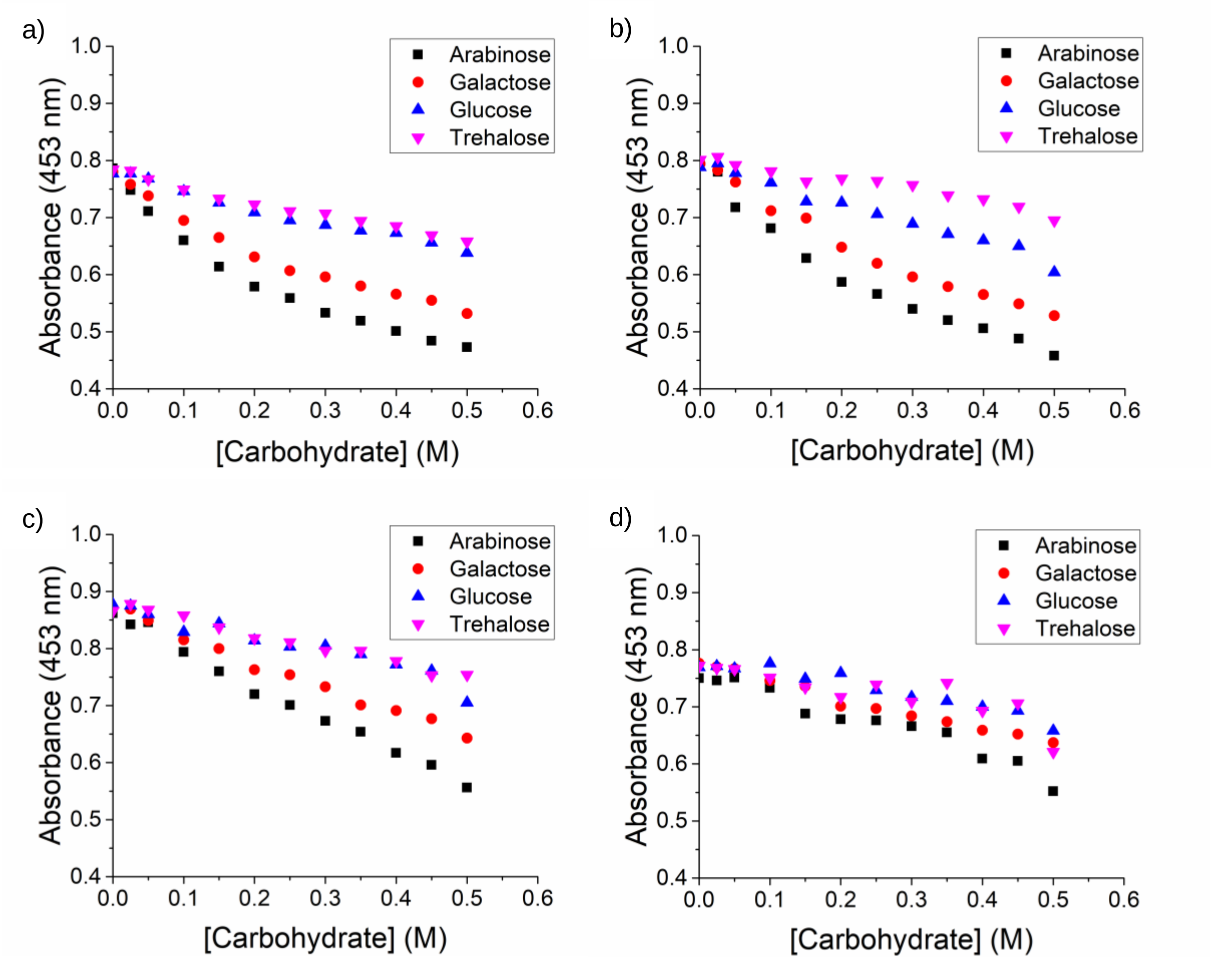


We were unable to test the binding of 2,2'-(ethylenedioxy)bis(ethylamine) G1 PAMAM tetra(3-phenylboronic acid)amide dendrimer **8** to the carbohydrates as it formed an insoluble complex with the alizarin red S dye.

**Figure S2. Structures of *Mtb* cell envelope components**. Trehalose monomycolate (TMM), trehalose dimycolate (TDM), phosphatidylinositol monomannoside (PIM 1), phosphatidylinositol monomannoside (PIM 2), phosphatidylinositol hexamannoside (PIM 6), phthiocerol dimycocerosate (PDIM)

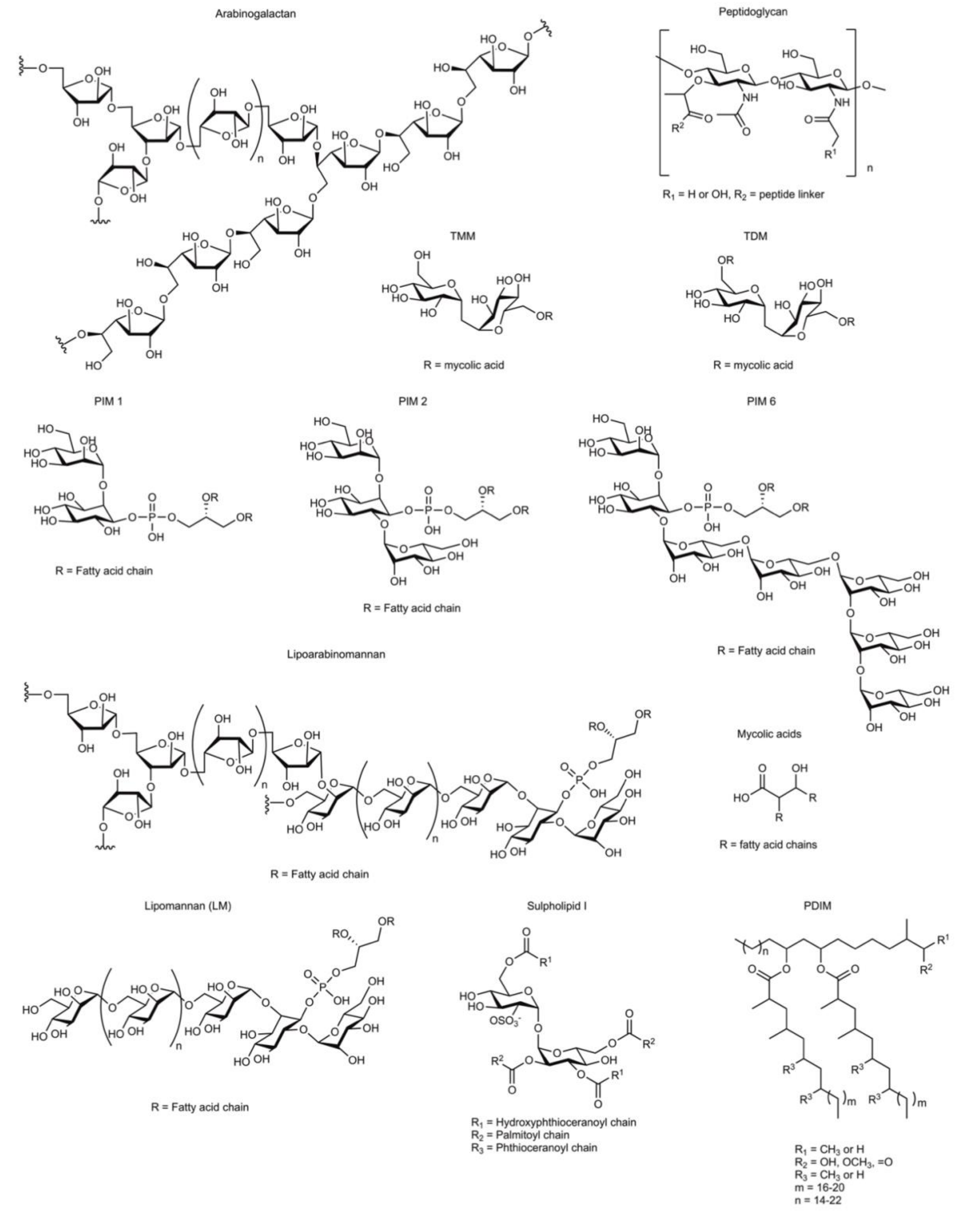

**Figure S3. Biolayer interferometry response to functionalised boronic acid sensors** a) Binding of isolated *Mtb* cell envelope components. The concentrations of the components are indicated (6.25 – 100 µg/mL). b) Binding to whole *E. coli* and gamma irradiated *Mtb* cells. The $OD_{600}$ are indicated (0 – 0.5). The synthesis and characterisation of the biotinylated boronic acid is described in Supplementary Methods and Results.

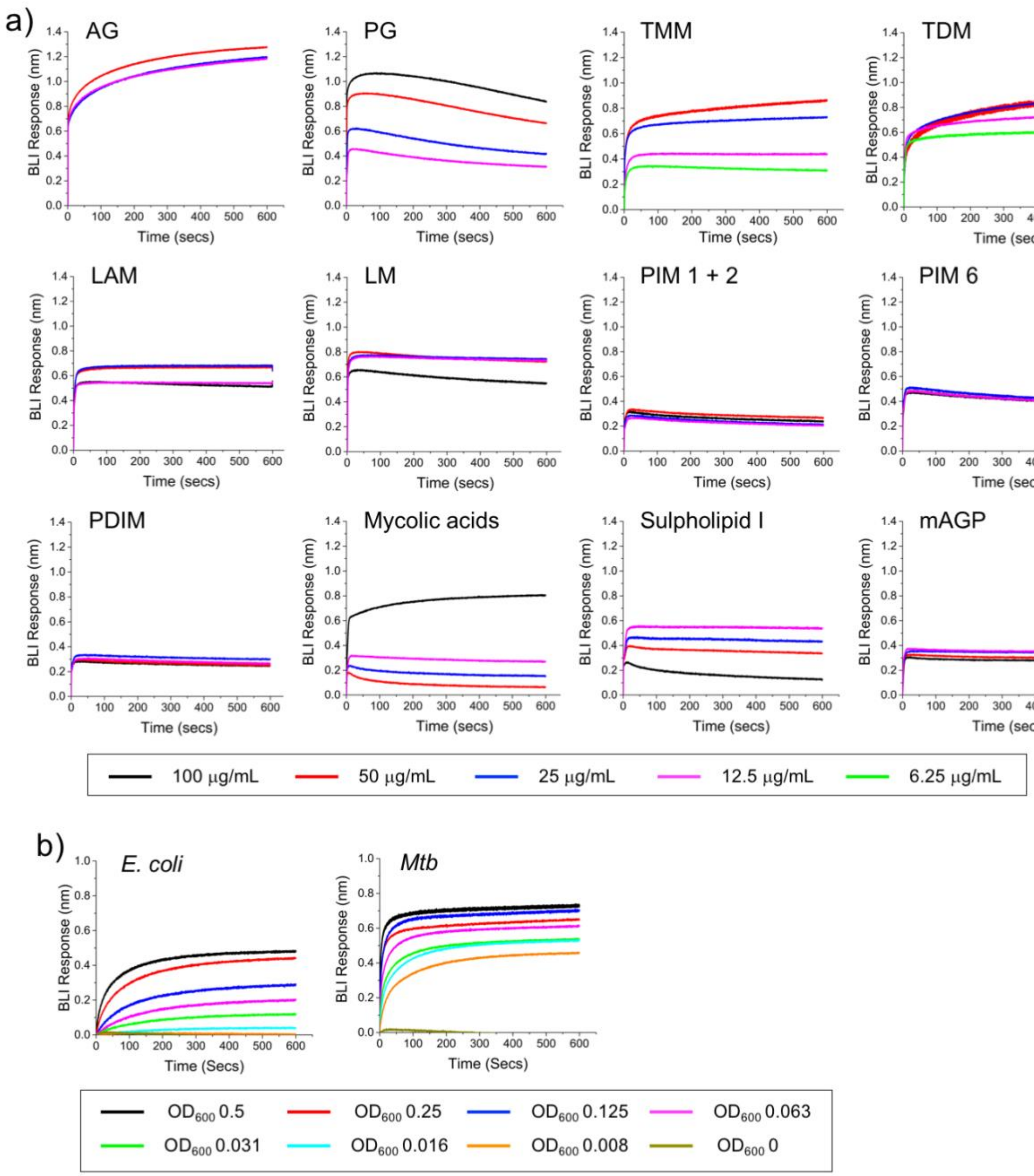

**Figure S4.** Biolayer interferometry response to functionalised boronic acid sensors to glycans not found in the mycobacterial cell wall. a) structure of dextran (from *Leuconostoc mesenteroides*) and galactan (from gum arabic). b) BLI response of functionalised sensors to dextran and galactan. The concentrations of the glycans are indicated.

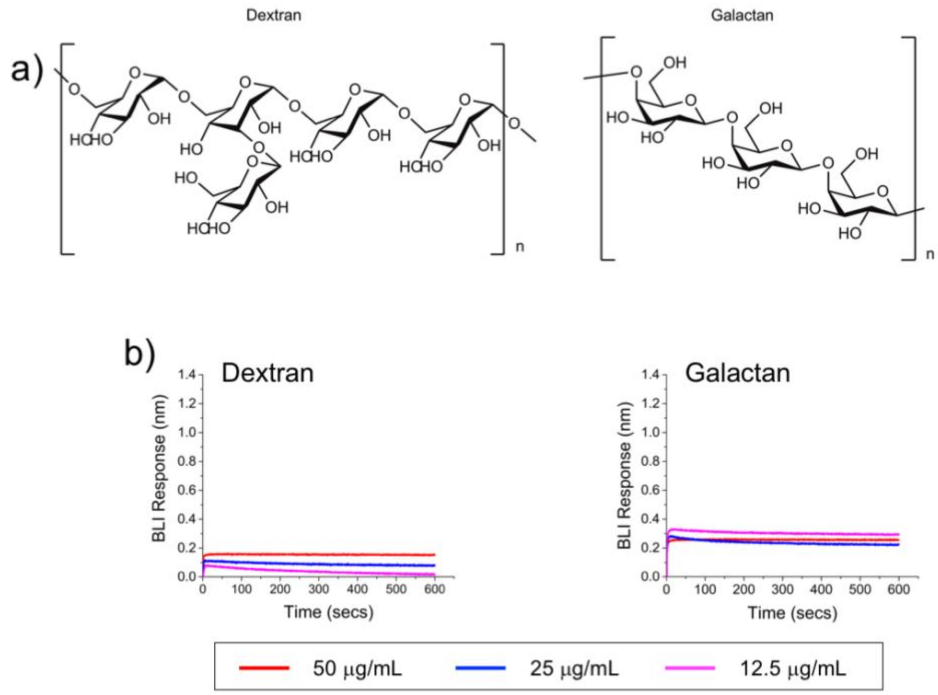

**Figure S5**. Representative images of checkerboard assays. a) checkerboard assay between compound **6** and meropenem showing additivity. b) Checkerboard assay between sulbactam (Sul) and meropenem (Mero) showing synergistic activity.

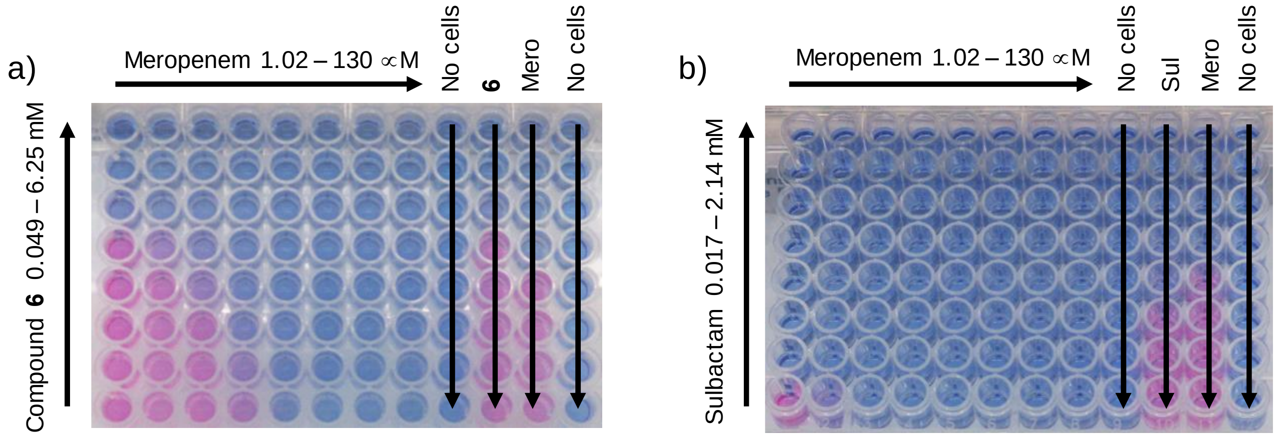

**Figure S6.** Functional distribution of total proteins identified by LC-MS in response to compound **6**. The functional categories are assigned according to Tuberculist.

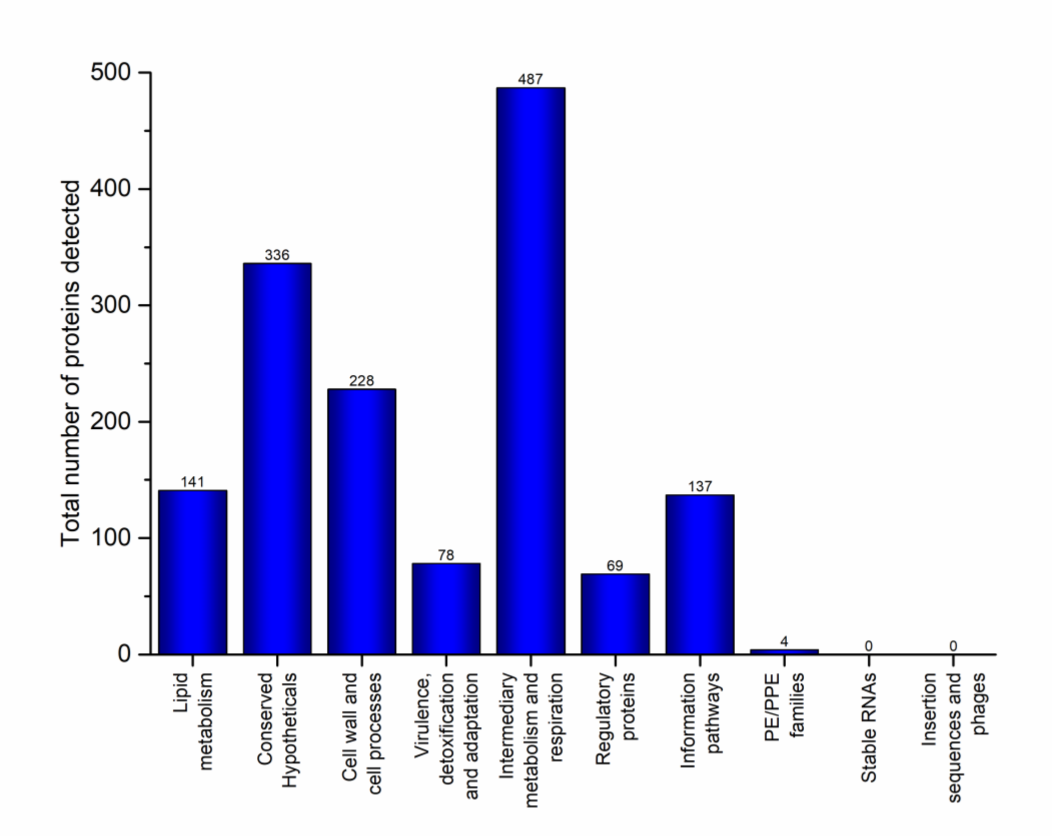

**Table S1**. Minimum bactericidal concentrations (MBC) of the dimeric boronic acids **4 - 6** against *Mycobacterium tuberculosis* H37Rv.

| **Compound** | **MBC** (mM) |
|---|---|
| **4** | 6.25 |
| **5** | 6.25 |
| **6** | 6.25 |

**Table S2**. Cytotoxicity testing of compounds **1-8** against human lung epithelium cell line and ovine blood.

| | **A549 ($MIC_{99}$) (a)** | **Red blood cells (b)** | |
|---|---|---|---|
| **Compound** | **mM** | **Haemolysis (%)** | **Agglutination** |
| **1** | >50 | 0 | ND |
| **2** | >25 | 4.5 | ND |
| **3** | >25 | 0.9 | ND |
| **4** | >50 | 0 | ND |
| **5** | >50 | 0 | ND |
| **6** | >50 | 0 | ND |
| **8** | >50 | 4.5(c) | ND |

a) A549 cells were exposed to compounds **1 - 8** for 24 h, and the cell viability was determined after this time by the addition of resazurin. The compounds were not toxic at the highest concentrations which are indicated. b) Haemolysis of ovine red blood cells in the presence of compounds **1 - 8**. Ovine red blood cells were exposed to compounds **1-8** for 1 hour after which the percentage haemolysis was determined by measuring the absorbance at 450 nm. Percentage lysis is compared with the 100% lysis after addition of lysis buffer (10 mM Tris, pH 7.8, 0.32 M sucrose, 5 mM $MgCl_2$, 10% Triton X-100). c) Due to solubility issues this compound was tested at 12.5 mM. ND – no agglutination detected.

**Table S3**. MIC values against *M. tuberculosis* H37Rv and corresponding interaction profiles with rifampicin and meropenem as determined by REMA checkerboard assays.

| **Compound** | **MIC (mM)** | **Interaction profile with Rifampicin** | | **Interaction profile with Meropenem** | |
|---|---|---|---|---|---|
| | | ∑FIC | Outcome | ∑FIC | Outcome |
| **4** | 3.13 | 2 | Additive | 1 | Additive |
| **5** | 3.13 | 2 | Additive | 0.625 | Additive |
| **6** | 1.56 | 2 | Additive | 0.625 | Additive |
| **Sulbactam** | 0.134 | - | - | 0.313 | Synergistic |

Fractional inhibitory concentrations (FICs) were calculated by use of the following formula as describe previously[1-3]: FIC ($X + Y$) = [MIC of compound $X$ in combination with $Y$]/[MIC of $X$ alone]. The fractional inhibitory index (ΣFIC) was calculated as FIC of compound $X$ + FIC of compound $Y$ to evaluate interaction profiles. ΣFICs of ≤0.5 designate synergistic activity, ΣFICs of ≥4.0 indicate antagonism, and values in between correspond to additivity.

### Supplementary Chemical and Synthetic protocols

#### General Information and Procedures

Unless stated, the chemicals and solvents, including anhydrous solvents, used in these syntheses were purchased from Sigma Aldrich and used as supplied and without further purification. Methanol (MeOH), dichloromethane (DCM), pyridine, triethylamine, ammonium hydroxide and magnesium sulfate ($MgSO_4$) were purchased from Fisher Scientific at laboratory reagent grade. Anhydrous tetrahydrofuran (THF) ≥99.9%, anhydrous *N,N*-dimethylformamide (DMF) ≥99.8%, anhydrous dichloromethane (DCM) ≥99.8%, anhydrous *N,N*-dimethylacetamide (DMAC) ≥99.8%, deuterium oxide ($D_2O$) 99.9%, oxalyl chloride 98%, ethylenediamine ≥99%, 2,2-(ethylenedioxy)bis(ethylenediamine) 98%, 4,7,10-trioxa-1,13-tridecanediamine 97% and methyl acrylate 99% were purchased from Sigma-Aldrich. 3-Carboxyphenylboronic acid 97%, 4-carboxyphenylboronic acid 97%, 3-aminophenylboronic acid hydrochloride 98% and 4-dimethylaminopyridine (DMAP) 99% were purchased from Acros Organics. Deuterochloroform ($CDCl_3$) 99.8% and deuteromethanol (MeOD) 99.8% were purchased from Apollo Scientific. Polyoxyethylene bis(amine) MW 1000 was purchased from Alfa Aesar. EZ-link Sulfo-NHS-LC-Biotin was purchased from Thermo Fisher. 1-(3-Dimethylaminopropyl)-3-ethylcarbodiimide HCl (EDCI) was purchased from Carbosynth. 100-500 MW dialysis tubing was obtained from Fisher Scientific. Distilled water ($H_2O$) was used throughout.

All reactions were performed using oven dried glassware and heat transfer was achieved using an oil bath.

Thin Layer Chromatography (TLC) was carried out using Merck aluminium backed sheets coated with 60 $F_{254}$ silica gel. Visualisation of the silica plates was achieved using a UV Lamp ($\lambda$ = 254 nm), and/or potassium permanganate (1.5 g $KMnO_4$, 10 g KCO3 and 1.25 mL 10% NaOH in 200 mL water) or Alizarin Red S (1 mM alizarin red S in acetone) [4]. Flash chromatography was carried out using Merck silica gel 60, 35-75 μm as the stationary phase (Sigma Aldrich). Mobile phases are reported in the ratio of solvents.

Proton ($^1H$) and carbon ($^{13}H$) NMR spectra were obtained at 298 K. $^1H$ NMR were recorded on Bruker DPX-300 and DPX-400 instruments as indicated. $^{13}C$ DEPT NMR were recorded on Bruker DPX-300 and DPX-400 instruments as indicated. NMR were fully assigned using COSY, HSQC and HMBC. All chemical shifts are quoted in parts per million (ppm), using the residual solvent as the internal standard. ($^1H$ NMR: $CDCl_3$ = 7.26, MeOD = 3.31, $D_2O$ = 4.79 and $^{13}C$ NMR: $CDCl_3$ = 77.0, MeOD = 49.0). Coupling constants (*J)* are reported in hertz (Hz) with the following abbreviations: s, singlet; d, doublet; t, triplet; q, quartet; quin, quintet; m, multiplet; br, broad.

Low resolution mass spectra (LRMS) were recorded on a Bruker Esquire 2000 spectrometer using electrospray ionisation (ESI) and high resolution mass spectra (HRMS) were recorded on either a Bruker HCT or Bruker HCT Ultra spectrometer. M/z values are reported in Daltons.

Infrared (IR) spectra were recorded on a Perkin-Elmer Avatar 320 FTIR spectrometer. Solids were compressed into a thin tablet and oils/liquids were analysed as films over a diamond sensor. Absorption maxima ($\upsilon_{max}$) are recorded in wavenumbers ($cm^{-1}$) and classified as strong (s) or broad (br).

Separation distances between boronic acid units were estimated assuming an extended chain with bond angles of 109° and bond lengths of 0.15 nm.

In addition to those specified above, the following abbreviations, designations and formulas are used throughout the supporting information: Aromatic (Ar), molecular weight (MW), polyamidoamine (PAMAM).

#### Synthetic procedures

##### ***N,N'*-(ethylenediamine)bis(3-phenylboronic acid)amide (4)**

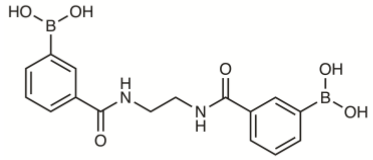


3-Carboxyphenylboronic acid (250 mg, 1.51 mmol) was dissolved in anhydrous THF (10 mL) under nitrogen and cooled to 0 °C. DMF (23 µL, 0.301 mmol) was then added, followed by the dropwise addition of oxalyl chloride (306 µL, 3.61 mmol). The reaction mixture was then allowed to warm to room temperature and stirred for two hours before removing the solvent *in vacuo*. The residue was then taken up in anhydrous DCM (10 mL) under nitrogen and cooled to 0 °C. Pyridine (122 µL, 1.51 mmol) and ethylenediamine (45 µL, 0.68 mmol) were added and the reaction mixture allowed to warm to room temperature and stirred for 16 hours. The solvent was then removed *in vacuo* and the product purified by column chromatography (5:1:0.1 DCM/MeOH/$H_2O$), to give the desired product **4** as a white solid (145 mg, 59%). $^1$H NMR (300MHz, MeOD) $\delta_{ppm}$ 8.63 (2H, br s, N<u>H</u>), 7.99 - 8.34 (2H, m, Ar H), 7.75-7.96 (4H, m, Ar H), 7.44 (2H, t, *J* = 7.5 Hz, Ar H), 3.64 (4H, m, CONHC<u>H</u>$_2$). $^{13}$C NMR (100MHz, $CDCl_3$) $\delta_{ppm}$ 169.7 (C=O), 136.6 (Ar <u>C</u>H), 133.3 (Ar <u>C</u>C=O), 132.3 (Ar <u>C</u>H), 128.6 (Ar <u>C</u>H), 127.4 (Ar <u>C</u>H), 39.6 (NH<u>C</u>H$_2$), (<u>C</u>B(OH$_2$) not observed). HRMS *m/z* (ES$^+$): [M+Na]$^+$ calcd. for $C_{16}H_{18}B_2N_2O_6Na^+$, 379.1243; found 379.1242. IR cm$^{-1}$: 3292 br (O-H), 2936 s (C-H), 1634 s (C=O).

##### ***N,N'*-(2,2-(ethylenedioxy)bis(ethylamine))bis(3-phenylboronic acid)amide (5)**

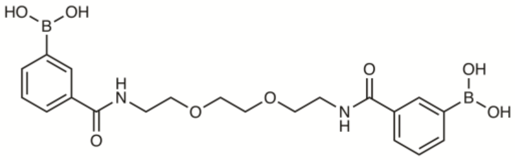


3-Carboxyphenylboronic acid (500 mg, 3.01 mmol) was dissolved in anhydrous THF (20 mL) under nitrogen and cooled to 0 °C. DMF (47 µL, 0.603 mmol) was then added, followed by the dropwise addition of oxalyl chloride (612 µL, 7.23 mmol). The reaction mixture was then allowed to warm to room temperature and stirred for two hours before removing the solvent *in vacuo*. The residue was then taken up in anhydrous DCM (20 mL) under nitrogen and cooled to 0 °C. Pyridine (244 µL, 3.01 mmol) and 2,2-(ethylenedioxy)bis(ethylamine) (198 µL, 1.36 mmol) were then added and the reaction mixture allowed to warm to room temperature and stirred for 16 hours. The solvent was then removed *in vacuo* and the product purified by column chromatography (5:1:0.1 DCM/MeOH/$H_2O$) to give the desired product **5** as a colourless oil (373 mg, 62%). $^1$H NMR (400MHz, MeOD) $\delta_{ppm}$ 8.37-8.61 (2H, m, N<u>H</u>), 7.97-7.27 (2H, m, Ar H)), 7.75-7.96 (4H, m, Ar H), 7.31-7.54 (2H, m Ar H), 3.63 –

3.72 (8H, m, $OC\underline{H}_2$), 3.53 – 3.61 (4H, m $CONHC\underline{H}_2$). $^{13}C$ NMR (100MHz, MeOD) $\delta_{ppm}$ 164.8 (C=O), 138.1 (Ar $\underline{C}$), 134.8 (Ar $\underline{C}C$=O), 133.7 (Ar $\underline{C}H$), 130.0 (Ar $\underline{C}H$), 128.8 (Ar $\underline{C}H$), 71.3 ($O\underline{C}H_2$), 70.6 ($O\underline{C}H_2$), 40.9 ($NH\underline{C}H_2$), ($\underline{C}B(OH_2)$ not observed). HRMS *m/z* ($ES^+$): $[M+Na]^+$ calcd. for $C_{20}H_{26}B_2N_2O_8Na^+$, 467.1767; found 467.1773. IR $cm^{-1}$: 3315 br (O-H), 2934 s (C-H), 1699 s (C=O).

***N,N'*-(4,7,10-trioxa-1,13-tridecanediamine)bis(3-phenylboronic acid)amide (6)**

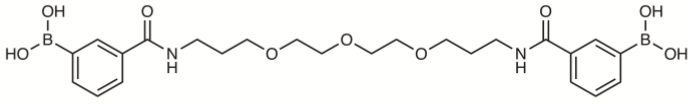

3-Carboxyphenylboronic acid (500 mg, 3.01 mmol) was dissolved in anhydrous THF (20 mL) under nitrogen and cooled to 0 °C. DMF (47 μL, 0.603 mmol) was then added, followed by the dropwise addition of oxalyl chloride (612 μL, 7.23 mmol). The reaction mixture was then allowed to warm to room temperature and stirred for two hours before removing the solvent *in vacuo*. The residue was then taken up in anhydrous DCM (20 mL) under nitrogen and cooled to 0 °C. Pyridine (244 μL, 3.01 mmol) and 4,7,10-trioxa-1,13-tridecanediamine (297 μL, 1.36 mmol) were then added and the reaction mixture allowed to warm to room temperature and stirred for 16 hours. The solvent was then removed *in vacuo* and the product purified by column chromatography (5:1:0.1 DCM/MeOH/$H_2O$) to give the product as a colourless oil (362 mg, 51 %). $^1H$ NMR (400MHz, MeOD) $\delta_{ppm}$ 8.16 (2H, m, Ar H), 7.78-7.94 (4H, m, Ar H), 7.43 (2H, t, *J* = 7.5 Hz, Ar H), 3.53 – 3.67 (12H, m, $OC\underline{H}_2$), 3.47 (4H, t, *J* = 6.5 Hz, $CONHC\underline{H}_2$), 1.87 (4H, quin, *J* = 6.0 Hz, $CH_2C\underline{H}_2CH_2$). $^{13}C$ NMR (100MHz, MeOD) $\delta_{ppm}$ 169.2 (C=O), 136.5 (Ar $\underline{C}H$), 133.6 (Ar $\underline{C}C$=O), 132.1 (Ar $\underline{C}H$), 128.5 (Ar $\underline{C}H$), 127.4 (Ar $\underline{C}H$), 70.1 ($O\underline{C}H_2$), 69.8 ($O\underline{C}H_2$), 69.0 ($O\underline{C}H_2$) 37.4 ($NH\underline{C}H_2$), 29.0 ($CH_2\underline{C}H_2CH_2$) ($\underline{C}B(OH_2)$ not observed). HRMS *m/z* ($ES^+$): $[M+Na]^+$ calcd. for $C_{24}H_{34}B_2N_2O_9Na^+$, 539.2348; found 539.2397. IR $cm^{-1}$: 3331 br (O-H), 2871 s (C-H), 1625 s (C=O).

***N,N'*-(polyoxyethylene bis(amine))bis(3-phenylboronic acid)amide (7)**

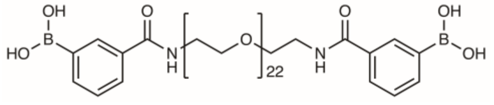

3-Carboxyphenylboronic acid (184 mg, 1.11 mmol), 1-ethyl-3-(3-dimethylaminopropyl)carbodiimide hydrochloride (320 mg, 1.67 mmol) and 4-(dimethylamino)pyridine (63 mg, 0.55 mmol) were dissolved in anhydrous DMF and heated to 60 °C for 30 mins. Polyoxyethylene bis(amine) MW 1000 (500 mg, 0.5 mmol) was then added and the reaction mixture heated to 60 °C for a further 16 hours. The reaction mixture was then cooled to room temperature and the solvent removed *in vacuo*. The residue was taken up in water (50 mL) and extracted with DCM (4 x 50 mL). The combined organic phases were dried ($MgSO_4$), filtered, and concentrated *in vacuo* to give the crude product as a brown oil. The oil was then dissolved in water (15 mL) and dialysed against water (100-500 MW dialysis tubing), and lyophilised to yield the desired product **7** as a brown solid (307 mg, 47%). $^1H$

NMR (400MHz, MeOD) $\delta_{ppm}$ 8.01 – 8.25 (2H, m, Ar H)), 7.61-8.01 (4H, m, Ar H), 7.26 – 7.52 (2H, m, Ar H), 3.49 – 3.84 (90H, m, OC$\underline{H}_2$), 3.39 -3.48 (4H, m, CONHC$\underline{H}_2$). $^{13}$C NMR (100MHz, $D_2O$) $\delta_{ppm}$ 171.5 ($\underline{C}$=O), 136.4 (Ar$\underline{C}$H), 132.7 (Ar$\underline{C}$C=O), 131.1 (Ar$\underline{C}$H), 131.1 (Ar$\underline{C}$H), 128.0 (Ar$\underline{C}$H), 71.7, 69.6, 68.9, 68.7 (O$\underline{C}H_2$), 39.5 (NH$\underline{C}H_2$). IR cm$^{-1}$: 3353 br (O-H), 2880 s (C-H), 1643 s (C=O).

**2,2'-(Ethylenedioxy)bis(ethylamine) G0.5 PAMAM dendrimer (9)**

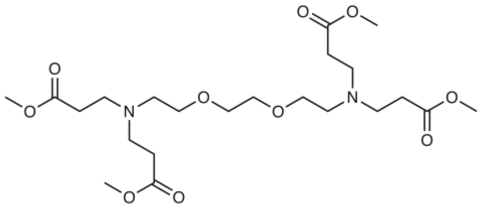

2,2'-(Ethylenedioxy)bis(ethylamine) (2 g, 13.5 mmol) was in dissolved in anhydrous MeOH (70 mL) under nitrogen and cooled to 0 °C. Methyl acrylate (6.08 mL, 67.5 mmol) was then added dropwise and the mixture stirred at room temperature for 96 hours. The reaction mixture was then cooled to room temperature and concentrated *in vacuo* at 40 °C to give the crude product as a pale yellow oil. The product was purified by column chromatography (19:1 DCM/MeOH to 9:1 DCM/MeOH) to give the desired product **9** as a pale yellow oil (4.24 g, 64 %). $^1$H NMR (400MHz, $D_2O$) $\delta_{ppm}$ 3.66 (12H, s, OC$\underline{H}_3$), 3.58 (4H, s, OC$\underline{H}_2$C$\underline{H}_2$O), 3.52 (4H, t, *J* = 6.0 Hz, OC$\underline{H}_2CH_2$N), 2.83 (8H, t, *J* =7.0 Hz, NC$\underline{H}_2CH_2$C=O), 2.67 (4H, t, *J* = 6.0 Hz, $OCH_2$C$\underline{H}_2$N), 2.46 (8H, t, *J* = 7.0 Hz, C=OC$\underline{H}_2CH_2$N). $^{13}$C NMR (100MHz, $D_2O$) $\delta_{ppm}$ 173.0 (C=O), 70.5 (O$\underline{C}H_2\underline{C}H_2$O), 69.6 (O$\underline{C}H_2CH_2$N), 53.2 ($OCH_2\underline{C}H_2$N), 51.5 (O$\underline{C}H_3$), 49.9 (N$\underline{C}H_2CH_2$C=O), 32.6 (C=O$\underline{C}H_2CH_2$N). HRMS *m/z* (ES$^+$): [M+Na]$^+$ calcd. for $C_{22}H_{41}N_2O_{10}^+$, 493.2756; found 493.2757. IR cm$^{-1}$: 2951 s (C-H), 2844 s (NC-H), 1730 s (C=O).

**2,2'-(Ethylenedioxy)bis(ethylamine) G1 PAMAM dendrimer (10)**

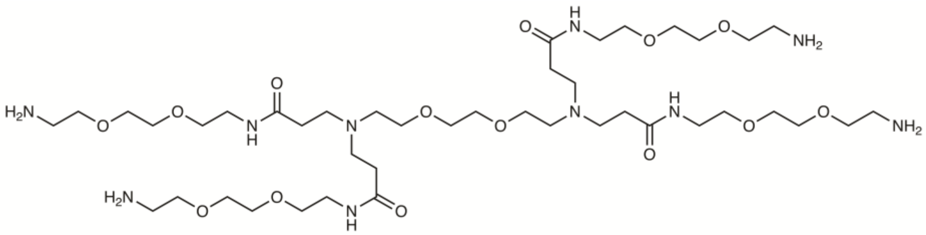

2,2'-(Ethylenedioxy)bis(ethylamine) G0.5 PAMAM dendrimer **9** **(**2g, 4.06 mmol) was dissolved in MeOH (5 mL) under nitrogen and cooled to -30 °C. 2,2'-(ethylenedioxy)bis(ethylamine) (47.4 mL, 325 mmol) was dissolved in MeOH (10 mL) under nitrogen and cooled to -30 °C. The solution of 2,2'-(Ethylenedioxy)bis(ethylamine) G0.5 PAMAM dendrimer was then added dropwise to the 2,2'-(ethylenedioxy)bis(ethylamine) solution under nitrogen, ensuring the temperature of the solution remained below -25 °C. The reaction was then slowly allowed to warm to room temperature and stirred for 9 days. The excess 2,2'-(ethylenedioxy)bis(ethylamine) was removed by vacuum distillation (117-120 °C at 10 mbar) to give the crude product as a pale yellow oil (3.75g). The product was then purified by column chromatography (20:9:1, DCM/MeOH/$NH_4OH$) to yield the desired product **10** as a pale yellow oil (1.7g, 43%). $^1H$ NMR (400MHz, $D_2O$) $\delta_{ppm}$ 7.68 (4H, br s, $CON\underline{H}$), 3.46 – 3.64 (40H, m, $OC\underline{H}_2$), 3.36 – 3.44 (8H, m, $CONHC\underline{H}_2$), 2.85 (8H, t, $J$ = 5.0 Hz, $C\underline{H}_2NH_2$), 2.77 (8H, t, $J$ = 6.0 Hz, $NC\underline{H}_2CH_2CO$), 2.66 (4H, t, $J$ = 5.5 Hz, $OCH_2C\underline{H}_2N(CH_2)CH_2$), 2.33 (8H, t, $J$ = 6.5 Hz, $C\underline{H}_2CONH$). $^{13}C$ NMR (100MHz, $D_2O$) $\delta_{ppm}$ 172.4 ($\underline{C}$=O), 73.2, 70.4, 70.2, 70.2, 69.9, 69.1 ($O\underline{C}H_2$), 52.8 ($OCH_2\underline{C}H_2N(CH_2)CH_2$), 50.5 ($N\underline{C}H_2CH_2CO$), 41.6 ($\underline{C}H_2NH_2$), 39.1 ($CONH\underline{C}H_2$), 33.90 ($\underline{C}H_2CONH$). HRMS *m/z* ($ES^+$): $[M+Na]^+$ calcd. for $C_{42}H_{88}N_{10}NaO_{14}^+$, 979.6374; found 979.6377. IR $cm^{-1}$: 3272 br (N-H), 2869 s (C-H), 1635 s (C=O).

**2,2'-(Ethylenedioxy)bis(ethylamine) G1 PAMAM tetra(3-phenylboronic acid)amide dendrimer (8)**

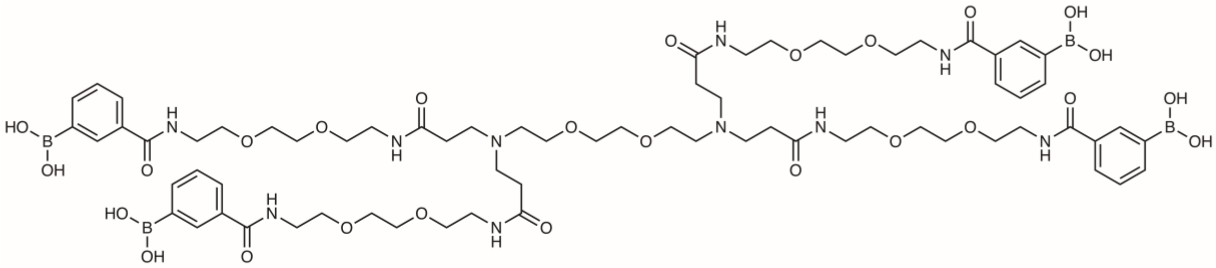

3-Carboxyphenylboronic acid (2.168 g, 13.1 mmol) was dissolved in anhydrous THF (60 mL) under nitrogen and cooled to 0 °C. DMF (202 μL, 2.6 mmol) was then added, followed by the dropwise addition of oxalyl chloride (2.65 mL, 31.4 mmol). The reaction mixture was then allowed to warm to room temperature and stirred for two hours before removing the solvent *in vacuo*. The residue was dissolved in anhydrous DMAC (40 mL) under nitrogen and cooled to 0 °C. Pyridine (1.05 mL, 13.1 mmol) and 2,2'-(Ethylenedioxy)bis(ethylamine) G1 PAMAM dendrimer **10** (0.5 g, 0.523 mmol) were dissolved in DMAC (20 mL) and added dropwise to the mixture at 0 °C. The reaction mixture allowed to warm to room temperature and stirred for 48 hours. The reaction mixture was then concentrated *in vacuo* and water (25 mL) was added and the reaction mixture then filtered. The filtrate was then dialysed against water (100-500 MW dialysis tubing) and lyophilised to yield the desired product as a yellow solid (465 mg, 57%). (NMR analysis showed the product **8** to be 95% tetrasubstituted).$^1H$ NMR (400MHz, $D_2O$) $\delta_{ppm}$ 7.93 (4H, s, $Ar\underline{H}$), 7.77 (4H, d, $J$ = 7.5 Hz, $Ar\underline{H}$), 7.68 (4H, d, $J$ = 7.5 Hz, $Ar\underline{H}$), 7.37 (4H, t, $J$ = 7.5 Hz, $Ar\underline{H}$), 3.14-3.74 (68H, m, $OC\underline{H}_2 + NC\underline{H}_2$), 2.56 (8H, t, $J$ = 5.5 Hz, $COC\underline{H}_2$).$^{13}C$ NMR (100MHz, $D_2O$) $\delta_{ppm}$ 171.5 ($\underline{C}$=O), 170.6 ($\underline{C}$=O), 137.2 ($Ar\underline{C}H$), 133.0 ($Ar\underline{C}C$=O), 132.1 ($Ar\underline{C}H$), 129.3 ($Ar\underline{C}H$), 128.3 ($Ar\underline{C}H$), 69.7 ($O\underline{C}H_2$), 69.5 ($O\underline{C}H_2$) 68.8 ($O\underline{C}H_2$), 68.6 ($O\underline{C}H_2$), 63.9 ($O\underline{C}H_2$), 53.1 ($OCH_2\underline{C}H_2N(CH_2)CH_2$), 50.1 ($N\underline{C}H_2CH_2CO$), 39.5

(CONH$\underline{C}H_2$), 38.9 (CONH$\underline{C}H_2$), 28.5 ($\underline{C}H_2$CONH), ($\underline{C}$B($OH_2$) not observed). HRMS *m/z* ($ES^+$): $[M+2H]^{2+}$ calcd. for $C_{70}H_{110}B_4N_{10}O_{26}^{2+}$, 775.3983; found 775.4093. IR $cm^{-1}$: 3276 (O-H) 2873 (C-H), 1634 (C=O).

**Biotinamido(hexanamido)(1,3-phenylboronic acid) (11)**

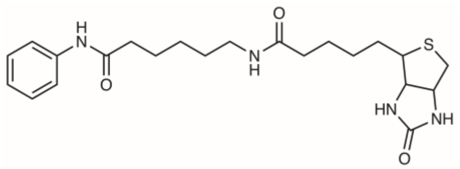

3-Aminophenylboronic acid hydrochloride (31 mg, 0.180 mmol) was dissolved in water (4 mL) and triethylamine (14 µL, 0.189 mmol) was added and the mixture stirred at room temperature for 10 mins. EZ-link Sulfo-NHS-LC-Biotin (100 mg, 0.180 mmol) was then added along with DMF (1 mL). The reaction mixture was stirred at room temperature for 16 hours and then concentrated *in vacuo*. The resulting brown oil was purified by column chromatography (4:1:0.1 DCM/MeOH/$H_2O$) to yield the desired product **9** as a pale brown oil (57 mg, 66%). $^1H$ NMR (400MHz, MeOD) $\delta_{ppm}$ 7.82 (1H, s, Ar $\underline{H}$), 7.65 (1H, d, *J* = 8.0 Hz, Ar $\underline{H}$), 7.51 (1H, d, *J* = 7.5 Hz, Ar $\underline{H}$), 7.32 (1H, t, *J* = 7.5 Hz, Ar $\underline{H}$), 4.47 – 4.56 (1H, m, $SCH_2C\underline{H}NH$), 4.27 – 4.37 (1H, m, SCHC$\underline{H}$NH), 3.13 – 3.27 (3H, m, SC$\underline{H}$ + CONHC$\underline{H}_2$), 2.88 – 2.99 (1H, m, SC$\underline{H}_aH_b$), 2.67 – 2.77 (1H, m, SC$\underline{H}_aH_b$), 2.41 (2H, t, *J* = 7.5 Hz, COC$\underline{H}_2$), 2.20 (2H, t, *J* = 7.5 Hz, COC$\underline{H}_2$), 1.33 – 1.81 (12 H, m, $CH_2C\underline{H}_2$). $^{13}C$ NMR (100MHz, MeOD) $\delta_{ppm}$ 174.91 (CNH$\underline{C}OCH_2$), 173.45 ($CH_2NH\underline{C}OCH_2$), 164.8 (NH$\underline{C}$ONH), 137.6 (Ar$\underline{C}$), 129.5 (Ar$\underline{C}$H), 127.8 (Ar$\underline{C}$H), 125.6 (Ar$\underline{C}$H), 122.3 (Ar$\underline{C}$H), 62.0 (SCH$\underline{C}$HNH), 60.2 ($SCH_2\underline{C}HNH$), 55.6 (S$\underline{C}$H), 39.7 (S$\underline{C}H_2$), 38.9 ($\underline{C}H_2$NH), 36.4 (CNHCO$\underline{C}H_2$), 35.5 ($CH_2NHCO\underline{C}H_2$), 28.7 ($\underline{C}H_2$), 28.3 ($\underline{C}H_2$), 28.0 ($\underline{C}H_2$), 26.1 ($\underline{C}H_2$), 25.5 ($\underline{C}H_2$), 25.2 ($\underline{C}H_2$). HRMS *m/z* ($ES^+$): $[M-H]^-$ calcd. for $C_{22}H_{32}BN_4O_5S^-$, 475.2192; found 475.2197.

#### Supplementary Biological protocols

##### Bacterial strains, cell lines, culture conditions and chemicals

*Mycobacterium smegmatis* $MC^2155$ (ATCC-700084) was routinely grown at 37 °C in Middlebrook 7H9 broth (BD Difco) supplemented with 0.2 % glycerol and 0.05 % Tween 80 or Tryptic Soy Broth (TSB) supplemented with 0.05 % Tween 80 or on Luria-Bertani (LB) agar. *Mycobacterium bovis* BCG (ATCC-35734) and *Mycobacterium tuberculosis* H37Rv were routinely grown at 37 °C in Middlebrook 7H9 broth supplemented with 0.2 % glycerol, 0.05 % Tween 80 and 10% albumin-dextrose-catalase (ADC) or on Middlebrook 7H10 plates supplemented with 0.5 % glycerol and 10% oleic acid-albumin-dextrose-catalase (OADC). *Escherichia coli* (Top 10) and *Pseudomonas putida* were routinely cultured in LB medium at 37 °C and 30 °C respectively. Human alveolar basal epithelial A549 cells (Public Health England, ECACC 86012804) were cultured at 37 °C with 5 % $CO_2$ atmosphere in Ham's F-12K (Kaighn's) Medium (Gibco, UK) supplemented with 10 % fetal-bovine serum, 100 Units/mL penicillin, 100 µg/mL streptomycin, and 250 ng/mL amphotericin B (HyClone). Ovine red blood cells were purchased from TCS Biosciences. PBST is phosphate buffered saline supplemented with 0.05% Tween 80.

All chemicals were purchased from Sigma-Aldrich unless otherwise stated. The galactan used was from gum arabic and was purchased from ACROS Organics. The dextran used was from *Leuconostoc mesenteroides*, with an average molecular weight of 150,000 and was purchased from Sigma-Aldrich.

##### *Mycobacterium tuberculosis* H37Rv reagents

The following reagents were obtained through BEI Resources, NIAID, NIH. All reagents are from *Mycobacterium tuberculosis* H37Rv*:* Gamma-Irradiated Whole Cells, NR-14819; Purified Lipoarabinomannan (LAM), NR-14848; Purified Lipomannan (LM), NR-14850; Purified Arabinogalactan, NR-14852; Purified Peptidoglycan, NR-14853; Purified Mycolylarabinogalactan-Peptidoglycan (mAGP), NR-14851; Purified Phthiocerol Dimycocerosate (PDIM), NR-20328; Purified Trehalose Monomycolate (TMM), NR-48784; Purified Trehalose Dimycolate (TDM), NR-14844; Purified Phosphatidylinositol Mannosides 1 & 2 (PIM1,2), NR-14846; Purified Phosphatidylinositol Mannoside 6 (PIM6), NR-14847; Purified Mycolic Acids, NR-14854; Purified Sulpholipid-1, NR-14845.

##### Alizarin Red S assay

Boronic acids **1-6** (3mM final concentration) were dissolved in alizarin red S (ARS) (0.144mM, in sodium phosphate buffer (0.1 M, pH 7.4)) (solution A) and added to the selected carbohydrate (0.5 M final concentration). The carbohydrate was serially diluted with solution A from 0-0.5M and the absorbance determined at 453 nm (BioTek Synergy HT Microplate Reader).

**Determination of minimum inhibitory concentrations**

The minimum inhibitory concentrations (MIC) of all compounds were determined using the resazurin reduction microplate assay (REMA) as described previously [5]. *M. smegmatis*, *M. bovis* BCG, *M. tuberculosis*, *E. coli and P. putida* were grown to mid-log phase and the inoculum standardized to 1 x $10^6$ colony forming units (CFU)/mL before addition to the prepared 96-well flat-bottom microtiter plate with 2-fold serial dilutions of each drug in media. An antibiotic control was also added to each plate (rifampicin for *M. smegmatis, M. bovis* BCG and *M. tuberculosis*, ampicillin for *E. coli* and tetracycline for *P. putida*) and the last column of the plate was used as a control without the addition of compound. The plates were incubated without shaking for 7 days (*M. bovis* BCG and *M. tuberculosis),* 24 hours (*M. smegmatis*), 16 hours (*E. coli* and *P. putida*) before addition of 25 µL resazurin (one tablet of resazurin (VWR) dissolved in 30 mL of sterile PBS). Following a further 24 hours incubation at 37 °C for mycobacteria or 2 hour incubation of *E. coli* and *P. putida* 37 °C and 30 °C respectively the plates were assessed for colour development. The MIC values were determined as the lowest concentration of drug that prevented the colour change of resazurin (blue – no bacterial growth) to resofurin (pink – bacterial growth).

**Determination of the minimum bactericidal (MBC) activity**

The MBC was determined for the compounds (**4 - 6**) against *M. tuberculosis* by setting up a microtiter plate as performed for the MIC determination. Instead of adding resazurin at 7 days each well from the microtiter plate was centrifuged (16,200 *x g*, 10 mins), supernatant discarded and the cells washed twice with PBS supplemented with 0.05% Tween 80 (PBST) before resuspending in PBST and plating on to Middlebrook 7H10 solid agar (supplemented with 0.5 % glycerol and 10% OADC). The plates were incubated at 37 °C for 3-4 weeks and the colony forming units (CFUs) determined. The lowest concentration at which no CFUs were observed was determined as the MBC.

**Cytotoxicity assay**

The cytotoxicity of the compounds was measured against a human lung epithelial cell line (A549). Briefly, cells were incubated ($10^6$ cells/well) with 2-fold serial dilutions of compounds in a 12 well plate (Thermo Fisher #150628), including a cell only control. Following incubation for 24 hours at 37 °C in the presence of 5 % $CO_2$ cell viability was determined by adding 100 µL resazurin (0.1% solution in PBS) for 1 h at 37 °C and measuring the absorbance of the resofurin metabolite at 570 nm (BioTek Synergy HT microplate reader). All assays were undertaken with 3 independent experimental repeats.

**Haemolysis assay**

The haemolysis activity of compounds **1-6** and **8** were tested against ovine red blood cells (TCS biosciences).

Samples were prepared by dissolving compounds in PBS, or PBS plus 20% DMSO for compounds **2**, **4**, **5** or PBS plus 33% DMSO compounds for compounds **3**, **6** and **8**. Two-fold serial dilutions were then carried out diluting with PBS. A positive control comprised lysis buffer (10 mM Tris, pH 7.8, 0.32 M sucrose, 5 mM $MgCl_2$, 10% Triton X-100), and the negative control comprised PBS (plus relevant % DMSO). 100 µL of ovine blood was incubated with 100 µL compound to give final concentrations of 0 - 50 mM for compounds **1-6** and 0 – 12.5 mM for compound **8** and samples incubated at room temperature for 1 hour, after which they were centrifuged for 5 min (2,000 *x g*, 22 °C). 20 µL of the supernatant was added to 750 µL of AHD solution (40 mM Triton X-100, 100 mM NaOH) and 200 µL was then added to 96 well plate and the absorbance read at 580 nm (BioTek Synergy HT Microplate Reader). The percentage lysis was calculated by comparison to the no compound control. All assays were undertaken with 3 independent experimental repeats.

**Agglutination assay**

The agglutination activity of compounds **1-6** and **8** were tested against ovine red blood cells. Samples were prepared by dissolving compounds in PBS, or PBS plus 20% DMSO for compounds **2**, **4**, **5** or PBS plus 33% DMSO compounds for compounds **3**, **6** and **8**. Two-fold serial dilutions were then carried out diluting with PBS. 100 µL of ovine blood was incubated with 100 µL compound to give final concentrations of 0 -50 mM for compounds **1-6** and 0 – 12.5 mM for compound **8**. Separately, 100 µL of 25% polyethylenimine was added as a positive control or 100 µL PBS (plus relevant % of DMSO) as a negative control. Following addition of the compounds, the ovine blood was incubated at room temperature for 1 hour and then the sample (50 µL) was added to a round-bottom 96-well microtiter plate (Corning, #3790) and incubated at room temperature for a further 30 min. The plate was then assessed for signs of agglutination (small red pellet at bottom of wells = no agglutination, red colouring across whole well = agglutination). All assays were undertaken with three independent experimental repeats.

**Determination of compound interactions using a REMA checkerboard assay**

A checkerboard assay was used to evaluate whether compound combinations act synergistically, antagonistically, or additively between compounds **4-6** with either rifampicin or meropenem against *M. tuberculosis* [1]. Compounds **4, 5** or **6** were serially diluted two-fold horizontally across the plate (0.049 – 6.25 mM final concentrations), and either rifampicin (5.93 – 759 nM final concentrations) or meropenem (1.02 – 130 µM final concentrations) was serially diluted two-fold vertically down the plate. Control wells in which compound **4**, **5** or **6** and rifampicin or meropenem alone were tested. *M. tuberculosis* was grown to mid-log phase and the inoculum standardised to 1 x $10^6$ colony forming units (CFU)/mL before addition to the plate. The plates were incubated for 7 days at 37 °C without shaking before addition of 25 µL resazurin (one tablet of resazurin (VWR) dissolved in 30 mL of sterile

PBS). Following a further 24 hours incubation at 37 °C the plates were assessed for colour change of the resazurin from blue (no bacterial growth) to pink (bacterial growth). Fractional inhibitory concentrations (FICs) were calculated by use of the following formula: FIC ($X + Y$) = [MIC of compound $X$ in combination with $Y$]/[MIC of $X$ alone]. The fractional inhibitory index (ΣFIC) was calculated as FIC of compound $X$ + FIC of compound $Y$ to evaluate interaction profiles. ΣFICs of ≤ 0.5 designate synergistic activity, ΣFICs of ≥ 4.0 indicate antagonism, and values in between correspond to additivity, as outlined in previous antibacterial combination studies [1-3].

**Resistance studies**

For single step resistance *M. bovis* BCG at $10^8$ CFUs were plated onto Middlebrook 7H10 agar supplemented with 0.5 % glycerol and 10% OADC containing 5 × MIC of compound **5**. A control in which no compound was added was also performed. The plates were then incubated at 37 °C for 3 months. No colonies were observed on the Middlebrook 7H10 agar plates that contained compound **5.**

**Biolayer Interferometry**

*E. coli* was grown to an $OD_{600}$ of 0.6. The cells were harvested (2,916 *x g*, 10 mins, 22 °C), washed three times with PBS and resuspended in PBS to a final $OD_{600}$ of 0.6 and serially diluted. Gamma-irradiated *M. tuberculosis* cells (BEI resources) were resuspended in PBS to give an $OD_{600}$ of 0.6 and serially diluted. Polysaccharides and cell wall components were dissolved in water (if necessary DMSO was used to dissolve the compound initially before diluting with water to give a maximum DMSO concentration of no greater than 5%) and serially diluted with water to give concentrations ranging from 0 - 100 μg/mL. Peptidoglycan and mycolylarabinogalactan-peptidoglycan (mAGP) did not fully dissolve and formed a suspension and were used at concentrations from 0 - 100 μg/mL. Biolayer Interferometry was carried out on ForteBio Octet Red96 (Forte Bio, USA). Assays were performed in black 96 well plates (Greiner Bio-one #655076) for assays with cells and black 96 well half area plates (Greiner bio-one #675076) for assays with polysaccharides and cell wall components. Assays were carried out at room temperature and PBS was used as the assay buffer. The wells were filled with 100 μL of either buffer or sample for assays with isolated cell wall components and polysaccharides and 200 μL of buffer or sample for assays with whole *E. coli* and *M. tuberculosis* cells and agitated at 1,000 rpm. Streptavidin (SA) biosensor tips (Forte Bio, USA) were hydrated in distilled water for at least 10 mins prior to use. The tips were functionalized by loading with the addition of 250 μg/mL biotinylated boronic acid **11** for 5 mins followed by a 5 mins wash step in PBS to remove unbound **11** and a stable baseline established. Following immobilization of the biotinylated boronic acid **11**, the binding interaction with different concentrations of isolated cell components, polysaccharides and whole cells was carried out which included baseline (5 min), association (10 min), dissociation (10 min). The $K_d$ values were calculated by plotting the end point association value against concentration (OriginPro 2016) and applying a logistic sigmodal fit then taking the EC50 as the $K_d$.

**Preparation of samples for proteomics**

*M. bovis* BCG was grown to an $OD_{600}$ of 0.3, and treated with 2x MIC compound **5** (3.125 mM). The cultures were incubated at 37 °C for 3 hours, 24 hours and 48 hours. At the time points indicated the cells were harvested (2,916 *x g*, 10 min, 22 °C), washed (2 x PBS), resuspended in SDC buffer (1% w/v sodium deoxycholate, 10 mM Tris(2-carboxyethyl)phosphine hydrochloride, 40 mM 2-chloroacetamide, 100 mM Tris pH 8.5) and heated to 95 °C for 10 mins. 0.1 mm silica glass beads were then added and the cells disrupted by bead-beating (4 x 45 secs on, 30 secs ice between cycles, Savant Bio 101 FastPrep FP120), followed by heating to 95 °C for 15 mins and sonication (water sonicator bath) at room temperature for 15 mins. The samples were centrifuged (16,200 *x g*, 10 mins) and the supernatant collected. The protein concentration was determined using the Qubit fluorometer (Thermo Fisher). Trypsin and LysC were added in 1:100 ratio of enzyme:protein and the samples digested at 37 °C overnight, followed by further purification through an Empore C18 solid phase extraction disk (3M, catalogue number 2215)

**NanoLC-ESI-MS/MS Analysis**

Protein Mass Spectrometry was performed on a Thermo Orbitrap Fusion (Thermo Scientific) coupled to an Ultimate 3000 RSLCnano HPLC (Dionex) using an Acclaim PepMap µ-precolumn cartridge (300 µm i.d. x 5 mm, 5 µm, 100 Å) and an analytical Acclaim PepMap RSLC column (75 µm i.d. x 50 cm, 2 µm, 100 Å, Thermo Scientific). Mobile phase buffer A was composed of 0.1% formic acid and mobile phase B was composed of acetonitrile containing 0.1% formic acid. The gradient was programmed as follows: 8% B increased to 25% B over 90 mins, then further increased to 35% B over 12 min, followed by 3 mins 90% B with a flow rate of 250 nL/min. Survey scans of peptide precursors from 375 to 1500 *m*/*z* were performed at 120K resolution (at 200 *m*/*z*) with a $2x10^5$ ion count target. The maximum injection time was set to 150 ms. Tandem MS was performed by isolation at 1.2 Th using the quadrupole, HCD fragmentation with normalized collision energy of 33, and rapid scan MS analysis in the ion trap. The $MS^2$ ion count target was set to $3x10^3$ and maximum injection time was 200 ms. Precursors with charge state 2–6 were selected and sampled for $MS^2$. The dynamic exclusion duration was set to 60 s with a 10 ppm tolerance around the selected precursor and its isotopes. Monoisotopic precursor selection was turned on and instrument was run in top speed mode.

**Data Analysis**

The raw data were searched using MaxQuant engine (V1.5.5.1) [6] against both the *Mycobacterium bovis BCG* and *Mycobacterium tuberculosis* databases and the common contaminant database from MaxQuant [7]. Peptides were generated from a tryptic digestion with up to two missed cleavages, carbamidomethylation of cysteines as fixed modifications, and oxidation of methionines as variable modifications. Precursor mass tolerance was 10 ppm and product ions were searched at 0.8 Da tolerances.

**Data processing and annotation**

Data processing and annotation were performed using the Perseus module of MaxQuant version 1.5.5.3 [6]. First, we eliminated the reverse and contaminant hits (as defined in MaxQuant) from the MaxQuant output files. Only protein groups identified with at least one uniquely assigned peptide and quantified with a minimum of two ratio counts were used for the analysis. For each experiment, the label free quantification intensity (LFQ) were transformed using the binary logarithm (log2). Protein groups were considered reproducibly quantified if identified and quantified in at least two replicates. Protein groups with significant relative regulation were determined according to the two sample test Student's T-Test with S0=0.5 and Permutation-based FDR 0.05. Protein categories were assigned based on Tuberculist [8] annotations.

#### References


1. Lechartier, B., Hartkoorn, R.C. & Cole, S.T. In Vitro Combination Studies of Benzothiazinone Lead Compound BTZ043 against *Mycobacterium tuberculosis. Antimicrob Agents and Chemother* **56**, 5790-5793 (2012).
2. Odds, F.C. Synergy, antagonism, and what the chequerboard puts between them. *J Antimicrob Chemother* **52**, 1 (2003).
3. Rand, K.H., Houck, H.J., Brown, P. & Bennett, D. Reproducibility of the microdilution checkerboard method for antibiotic synergy. *Antimicrob Agents Chemother* **37**, 613-615 (1993).
4. Duval, F., van Beek, T.A. & Zuilhof, H. Sensitive Thin-Layer Chromatography Detection of Boronic Acids Using Alizarin. *Synlett*, 1751-1754 (2012).
5. Palomino, J.C. et al. Resazurin microtiter assay plate: Simple and inexpensive method for detection of drug resistance in *Mycobacterium tuberculosis*. *Antimicrob Agents Chemother* **46**, 2720-2722 (2002).
6. Cox, J. & Mann, M. MaxQuant enables high peptide identification rates, individualized p.p.b.-range mass accuracies and proteome-wide protein quantification. *Nat Biotechnol* **26**, 1367-72 (2008).
7. Tyanova, S., Temu, T. & Cox, J. The MaxQuant computational platform for mass spectrometry-based shotgun proteomics. *Nat Protoc* **11**, 2301-2319 (2016).
8. Kapopoulou, A., Lew, J.M. & Cole, S.T. The MycoBrowser portal: a comprehensive and manually annotated resource for mycobacterial genomes. *Tuberculosis (Edinb)* **91**, 8-13 (2011).

#### NMR spectra

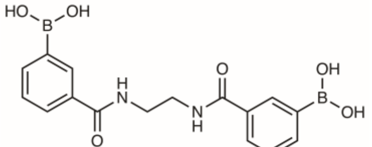


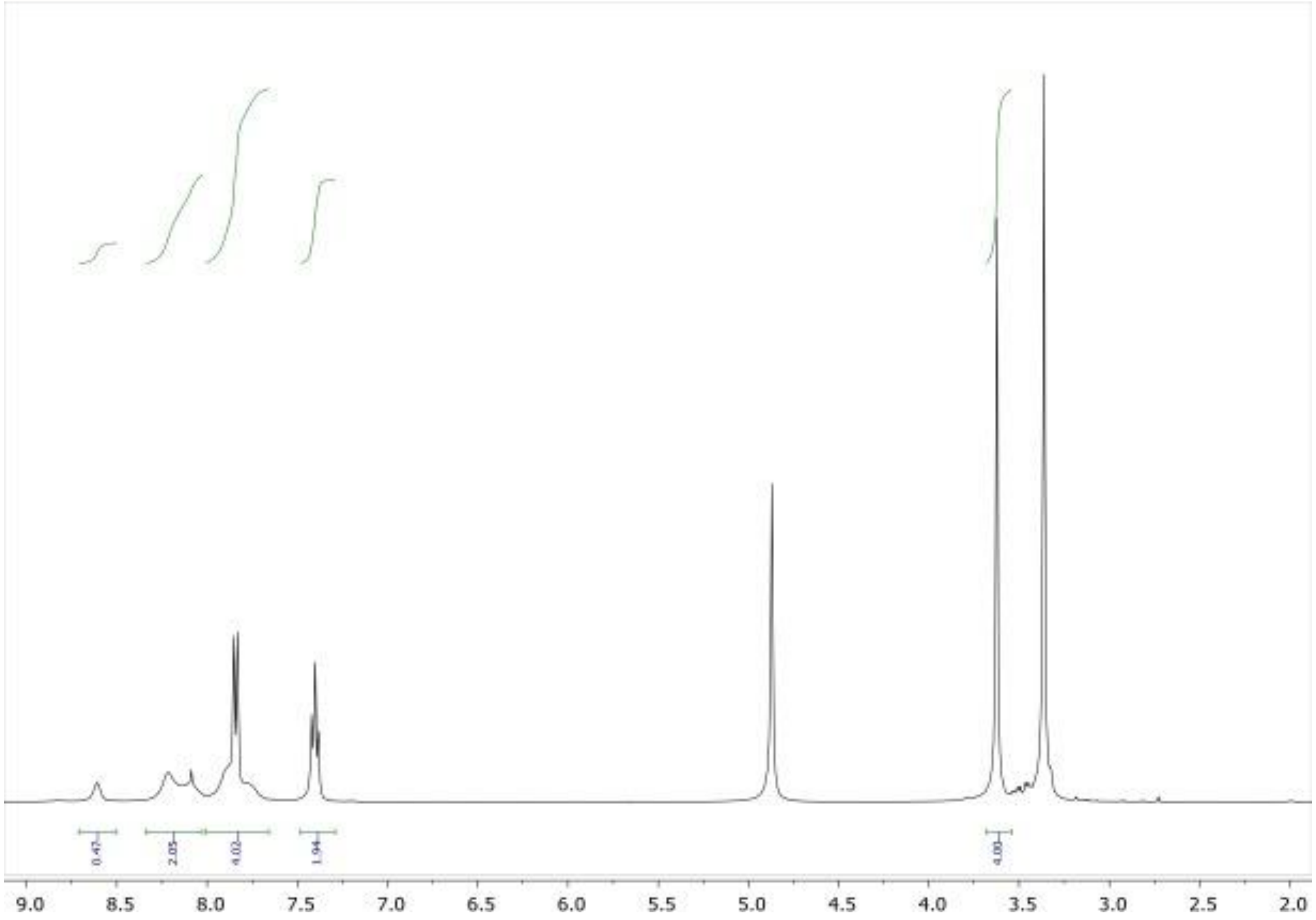


**Figure S7**. $^{1}$H NMR spectrum of compound **4**

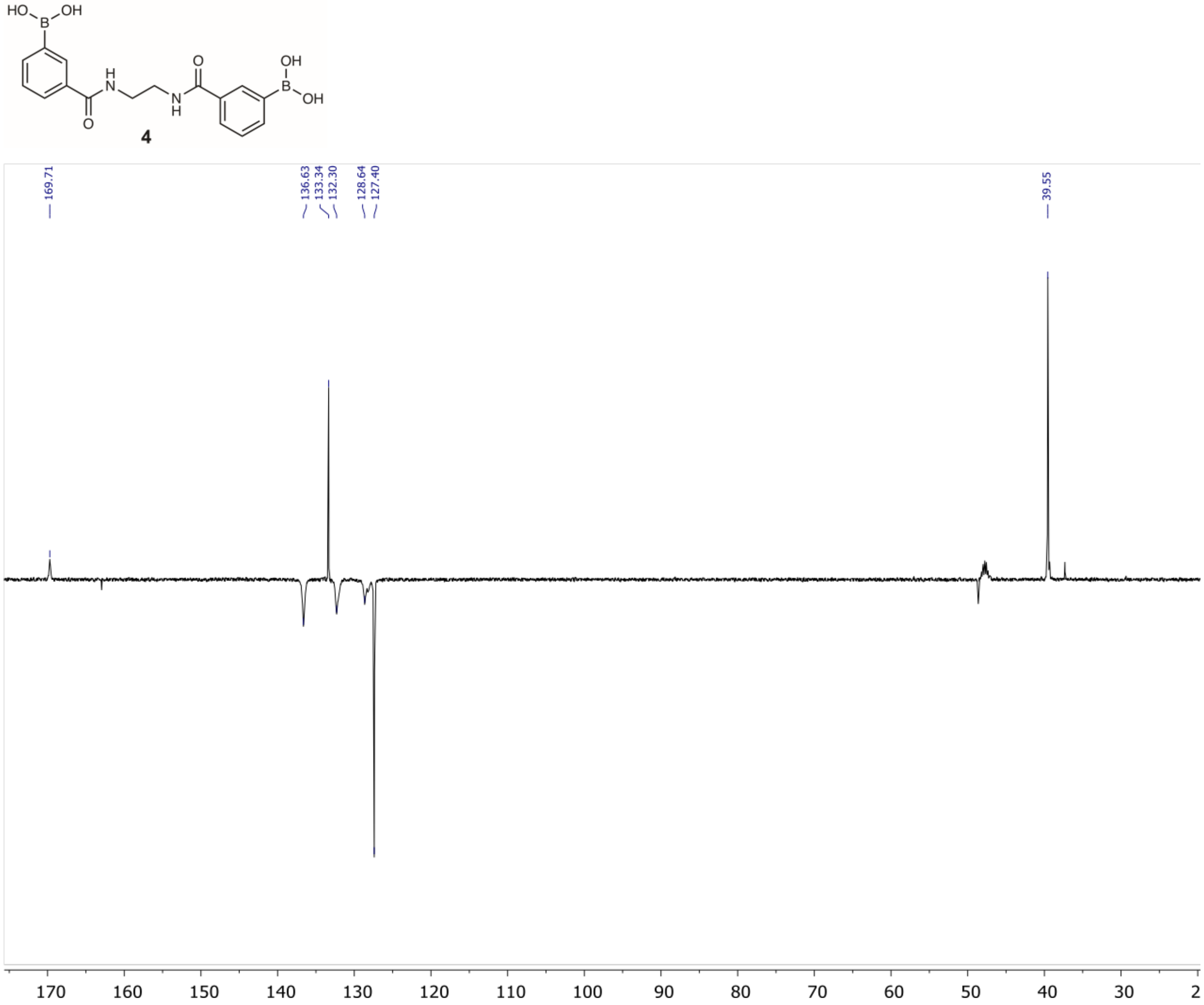


**Figure S8**. $^{13}$C DEPT NMR spectrum of compound **4**

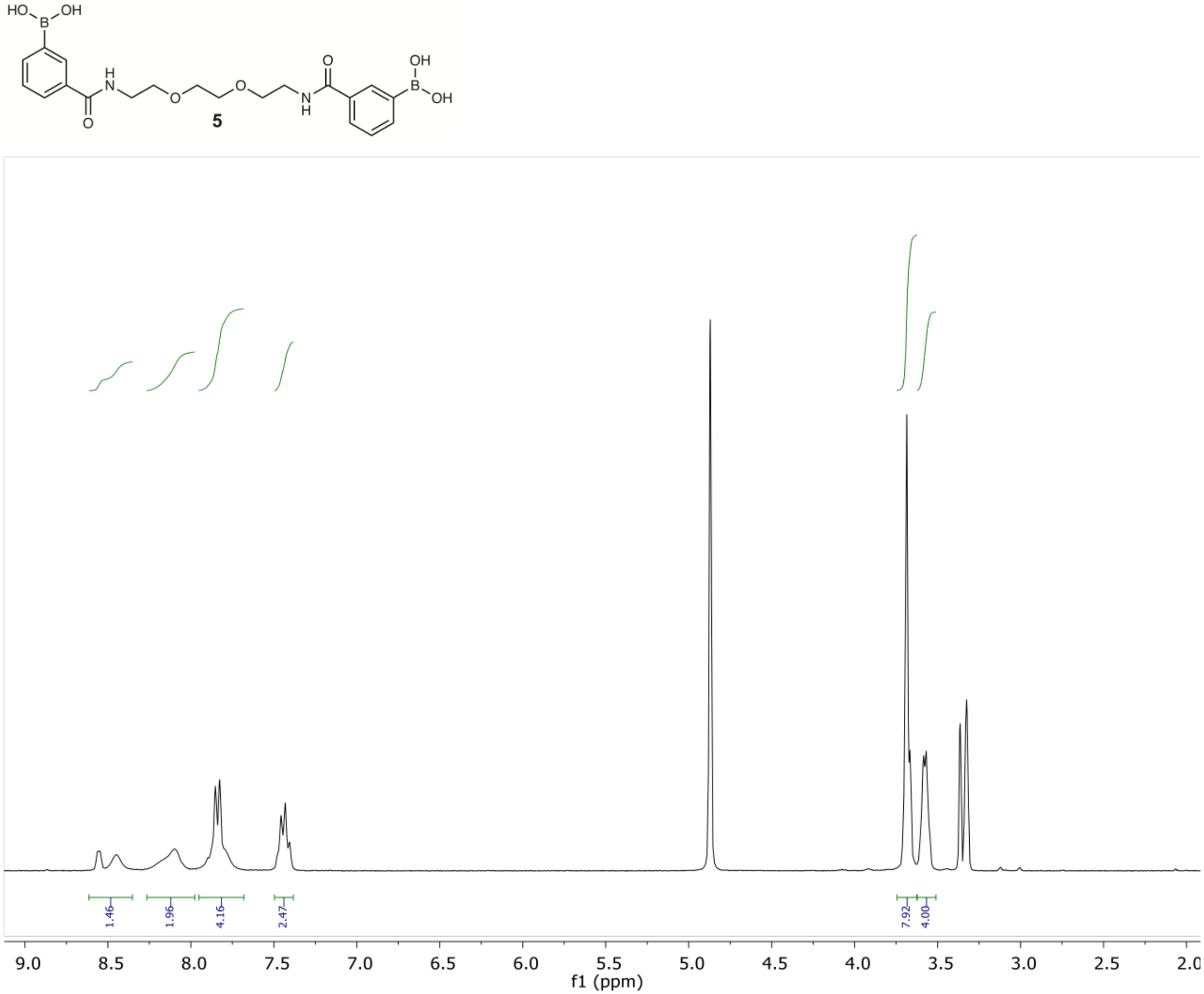


**Figure S9**. $^{1}$H NMR spectrum of compound **5**

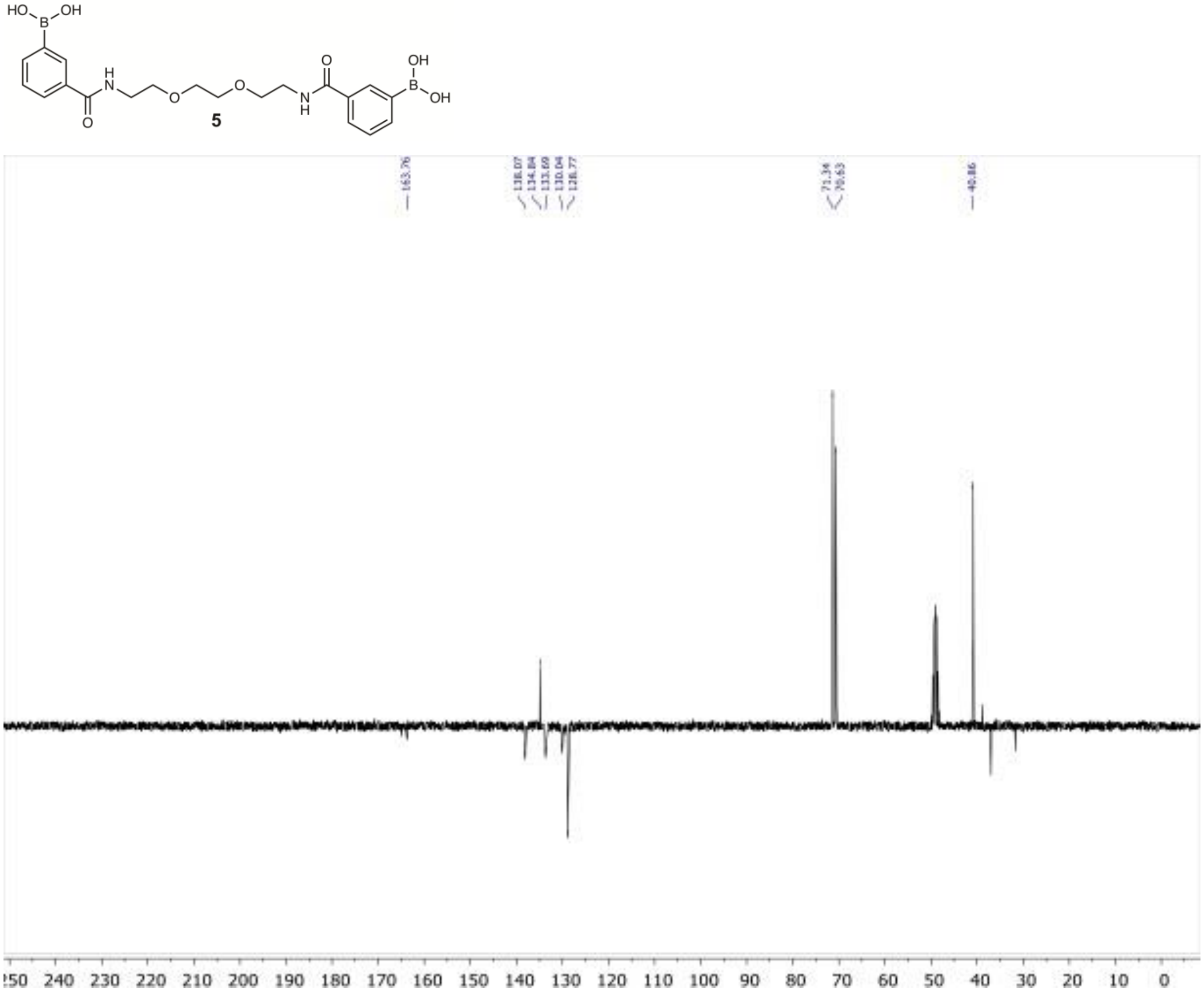


**Figure S10**. $^{13}$C DEPT NMR spectrum of compound **5**

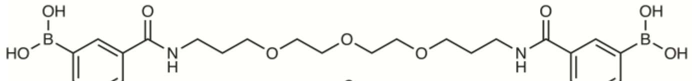


**6**

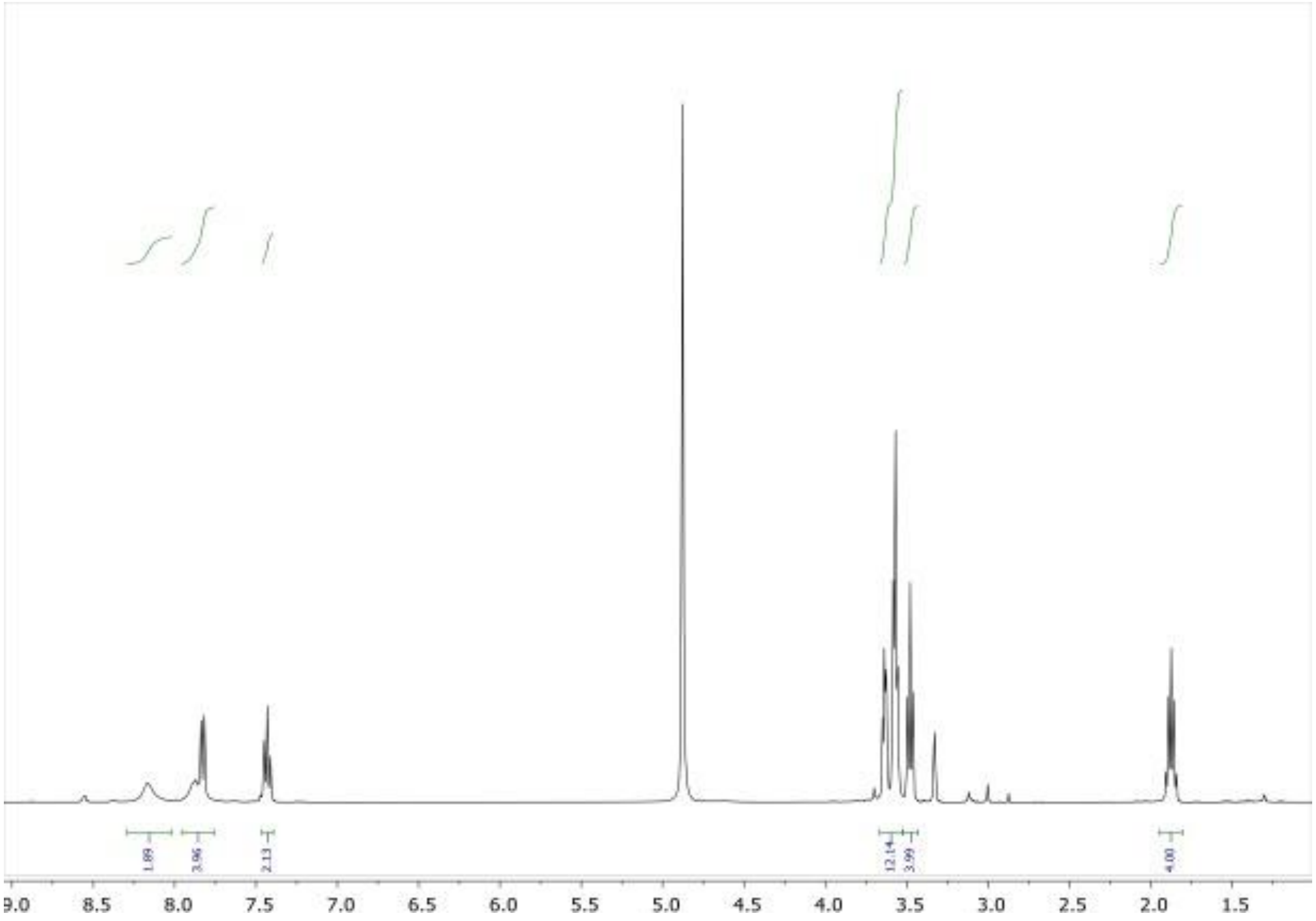


**Figure S11**. $^{1}$H NMR spectrum of compound **6**

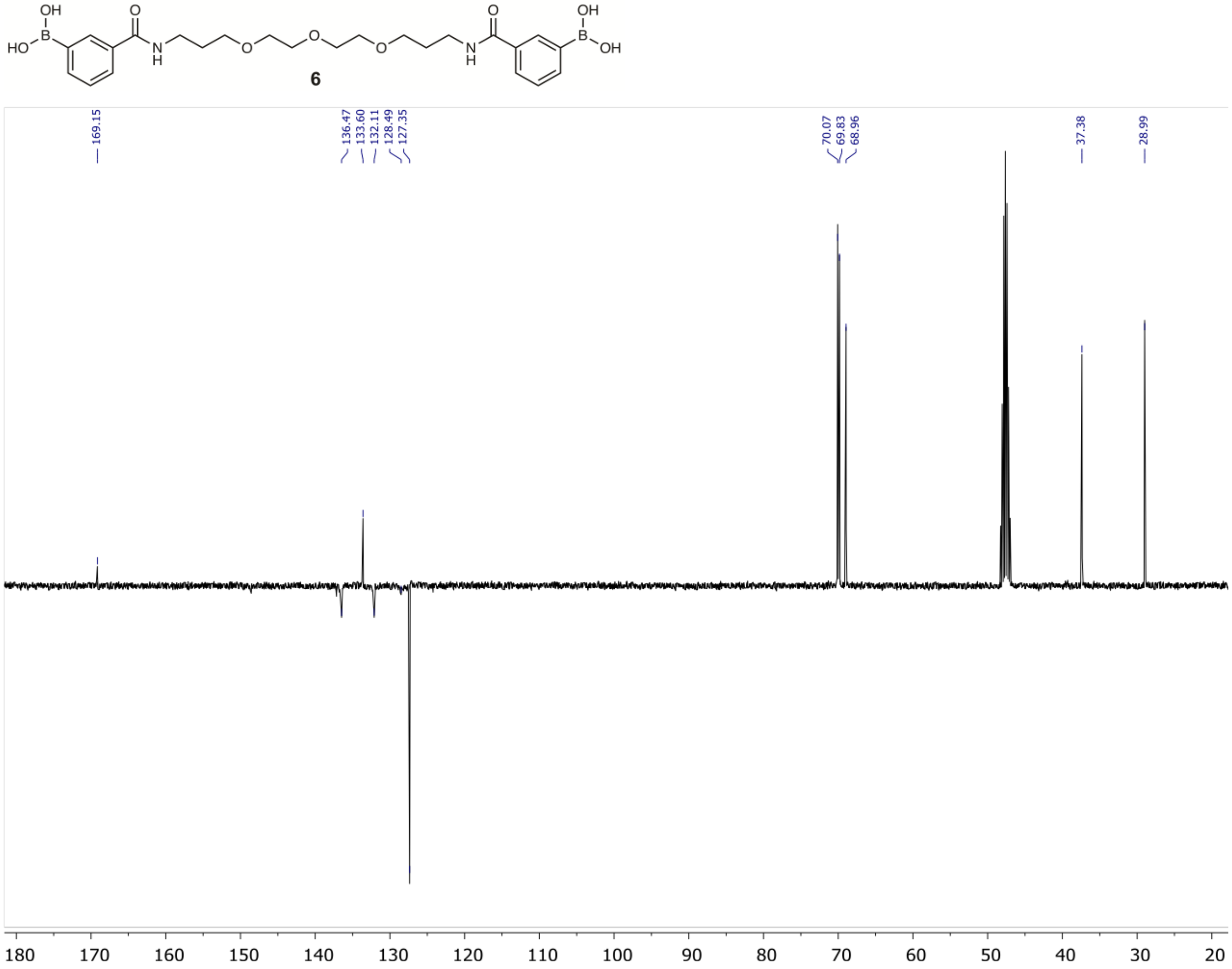


**Figure S12**. $^{13}C$ DEPT NMR spectrum of compound **6**

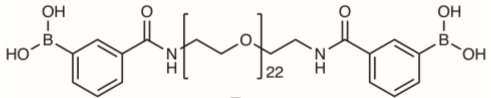

**7**

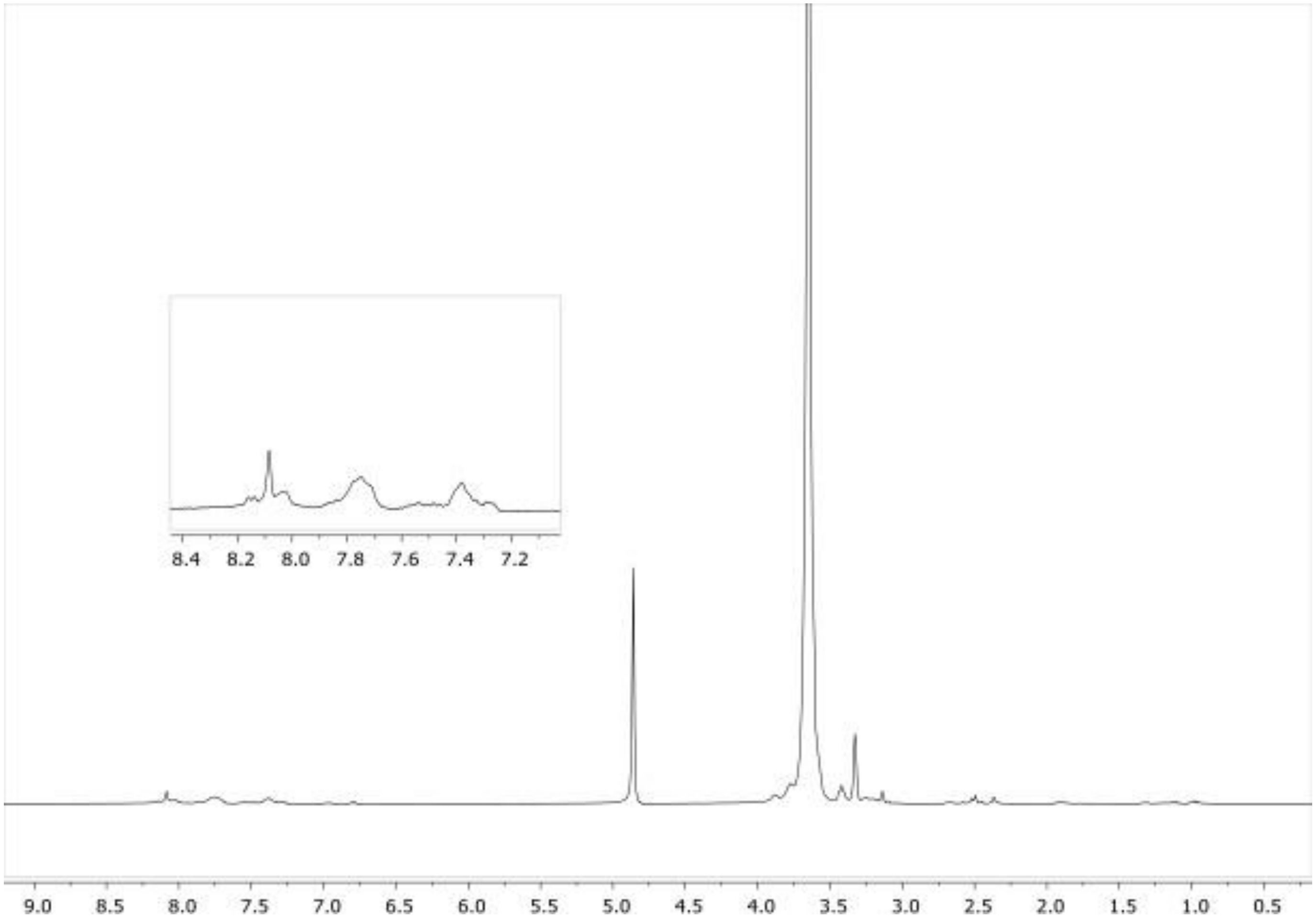


**Figure S13**. $^{1}$H NMR spectrum of compound **7**

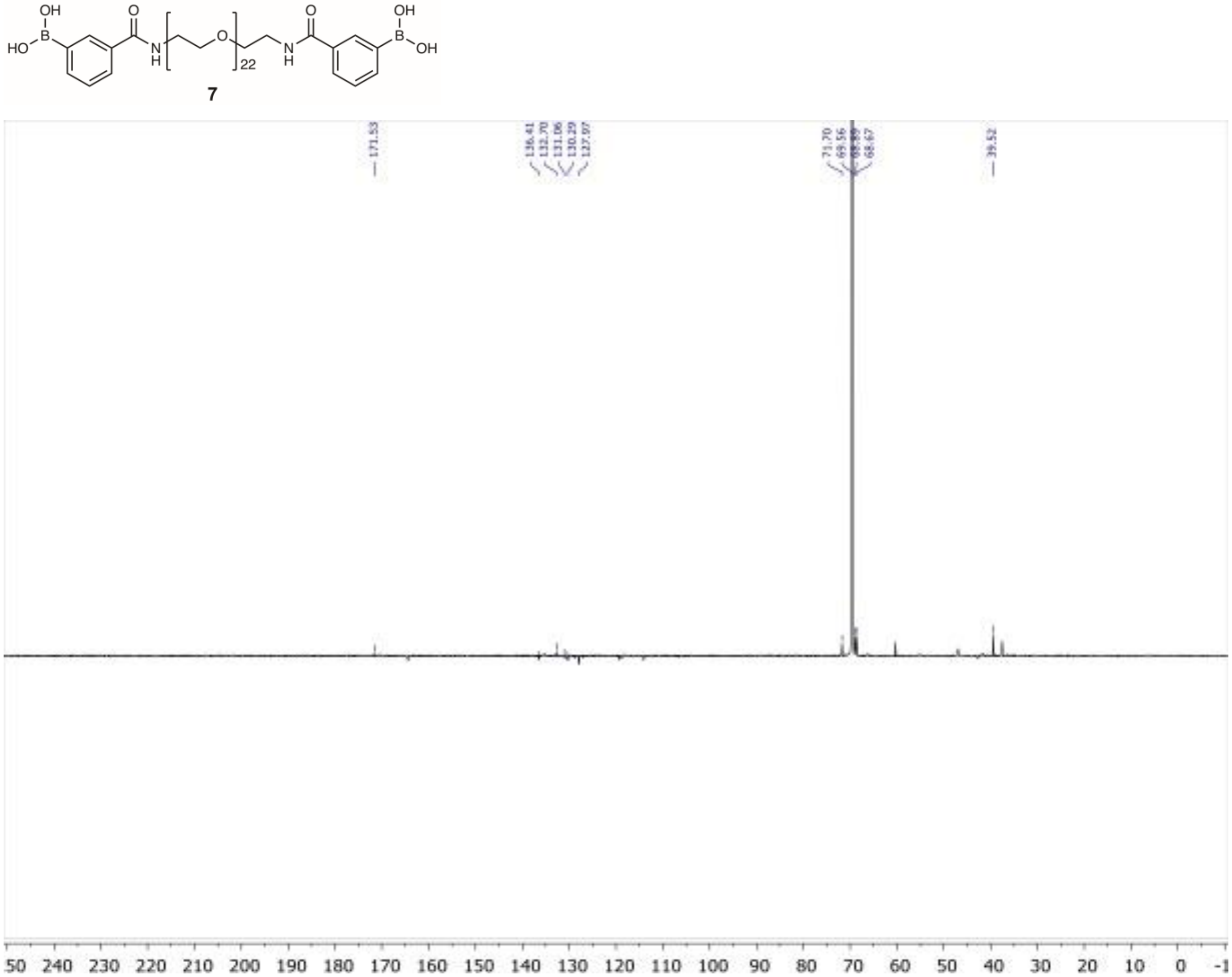


**Figure S14**. $^{13}$C DEPT NMR spectrum of compound **7**

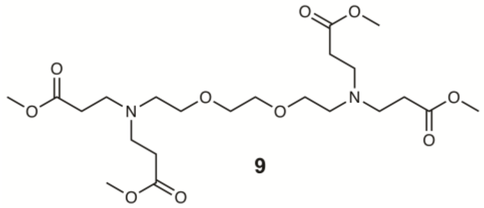

9

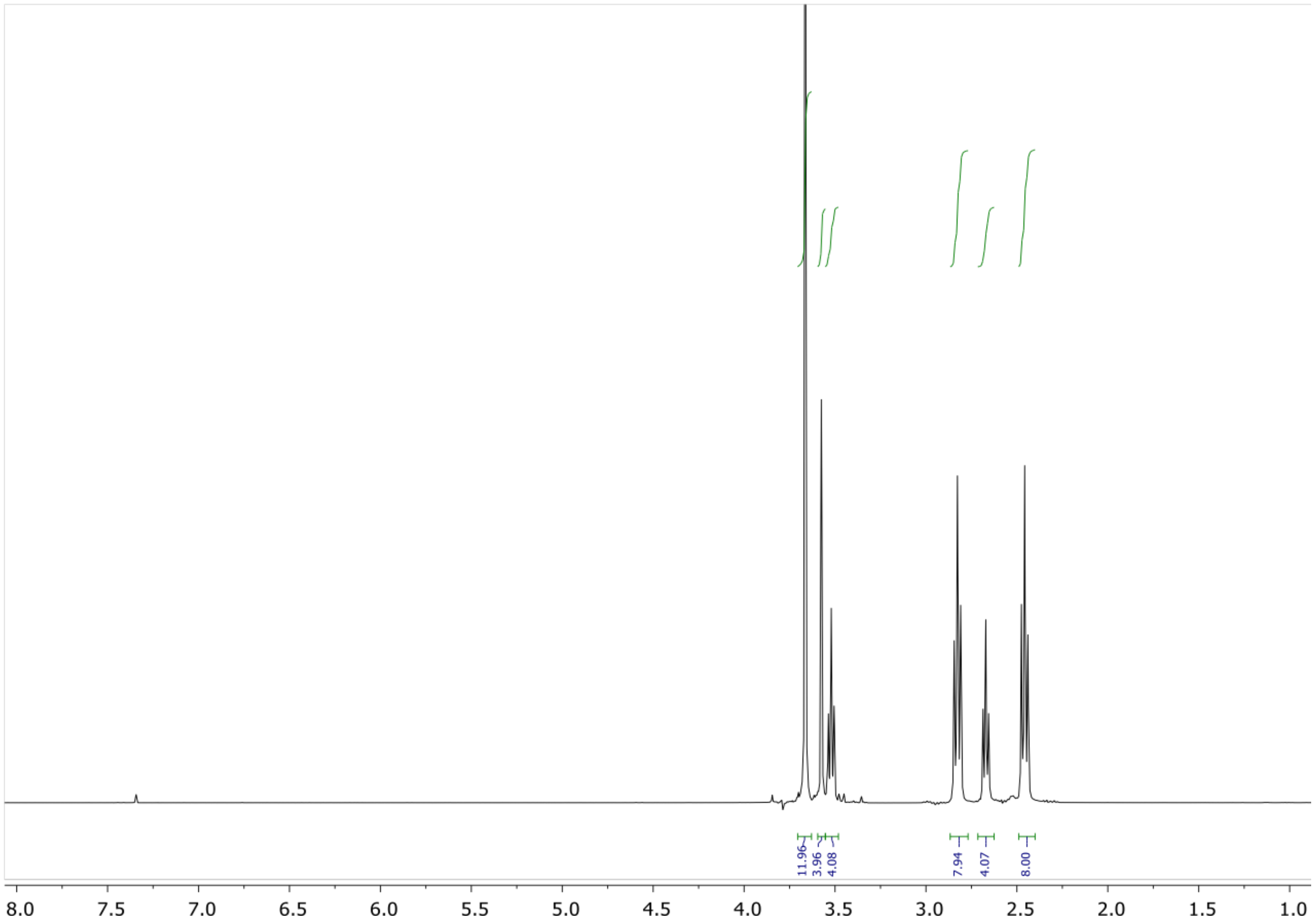


**Supplementary Figure S15**. $^{1}$H NMR spectrum of compound **9**

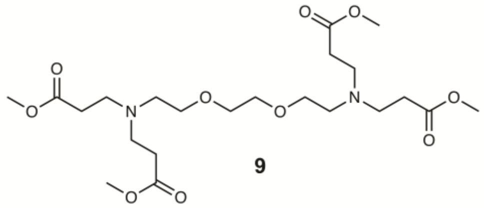

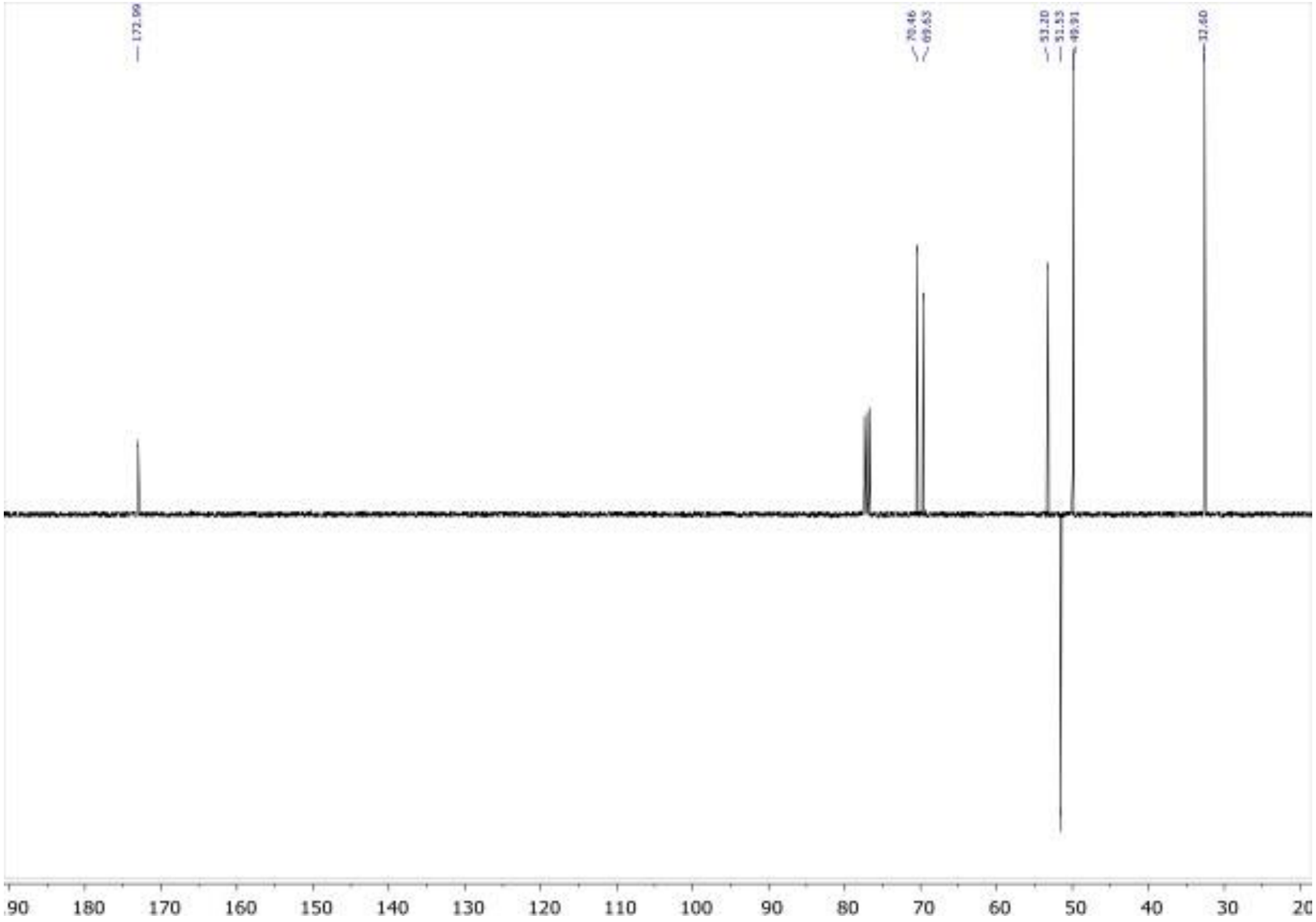


**Figure S16**. $^{13}$C DEPT NMR spectrum of compound **9**

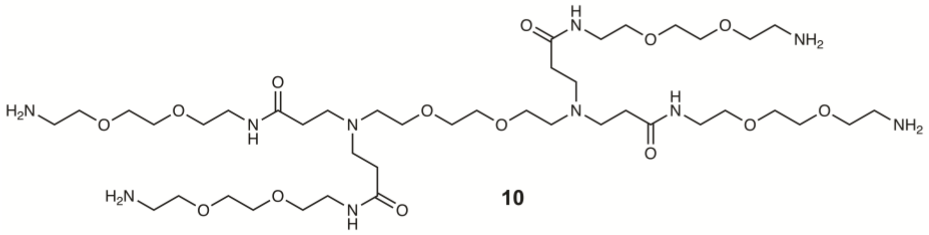

**10**

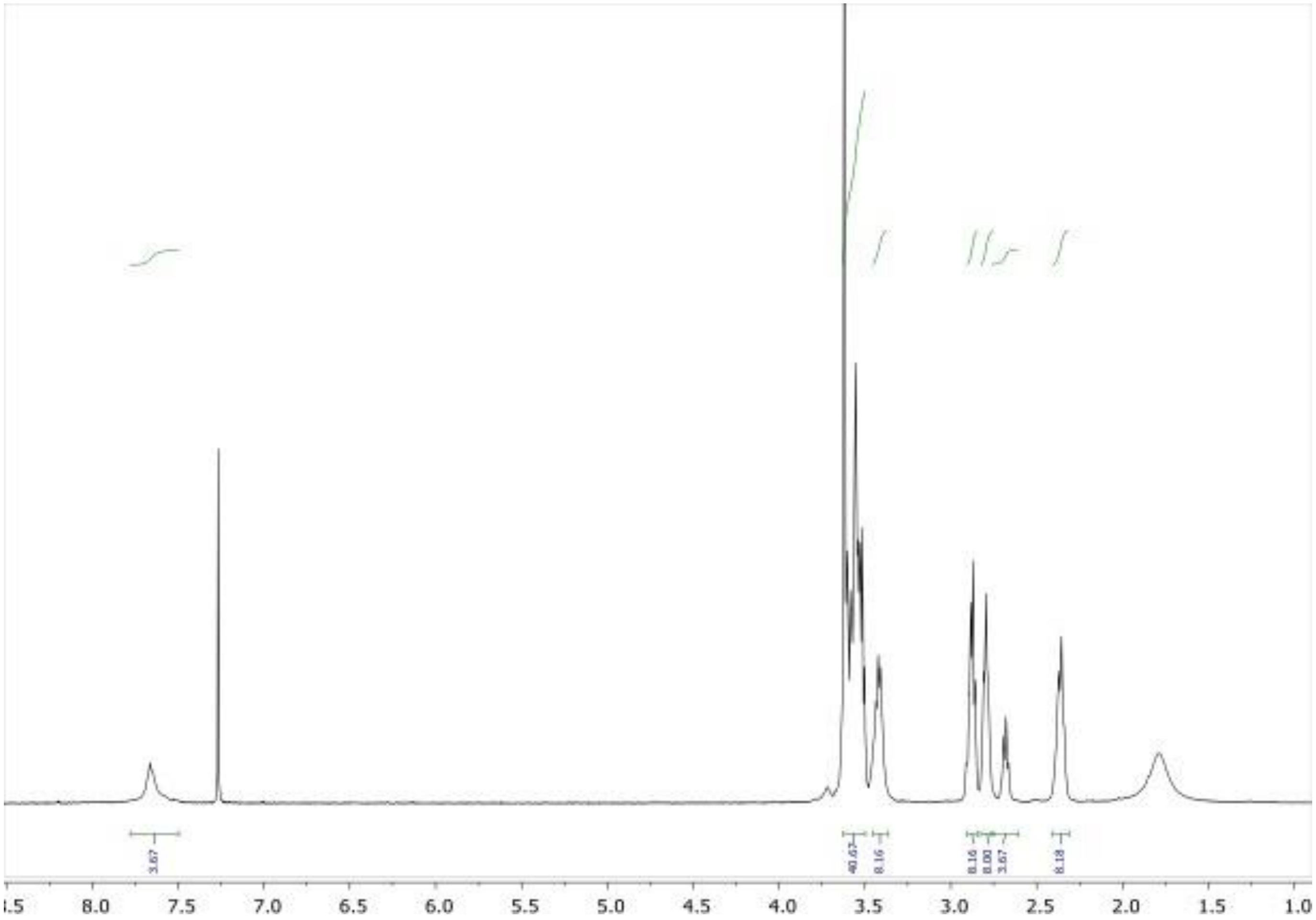


**Figure S17**. $^{1}H$ NMR spectrum of compound **10**

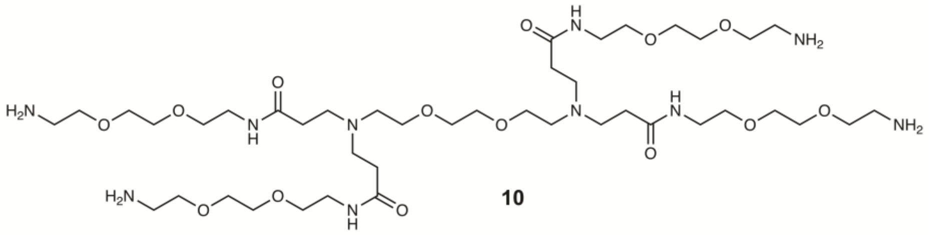

**10**

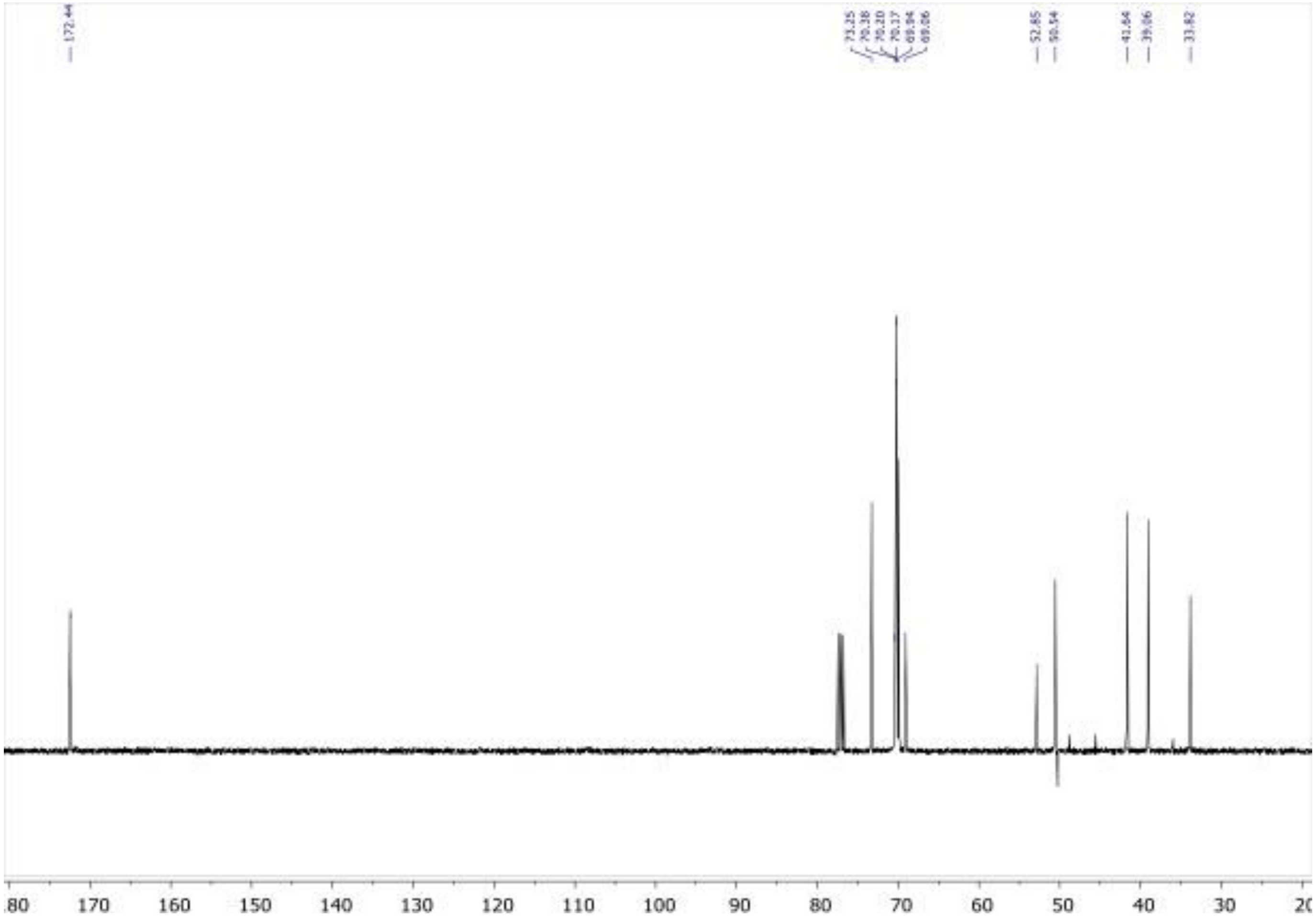


**Figure S18**. $^{13}$C DEPT NMR spectrum of compound **10**

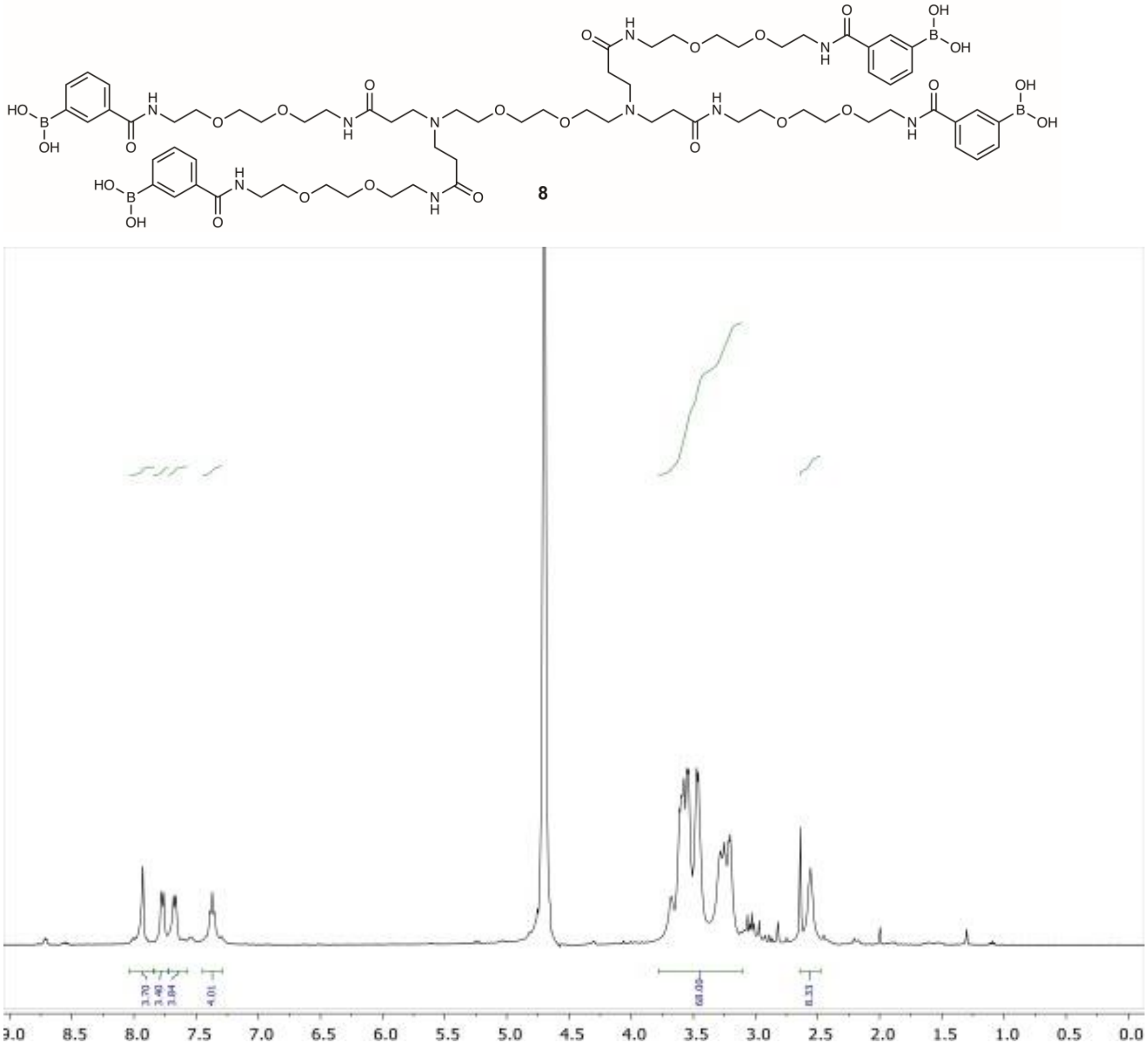


**Figure S19**. $^{1}$H NMR spectrum of compound **8**

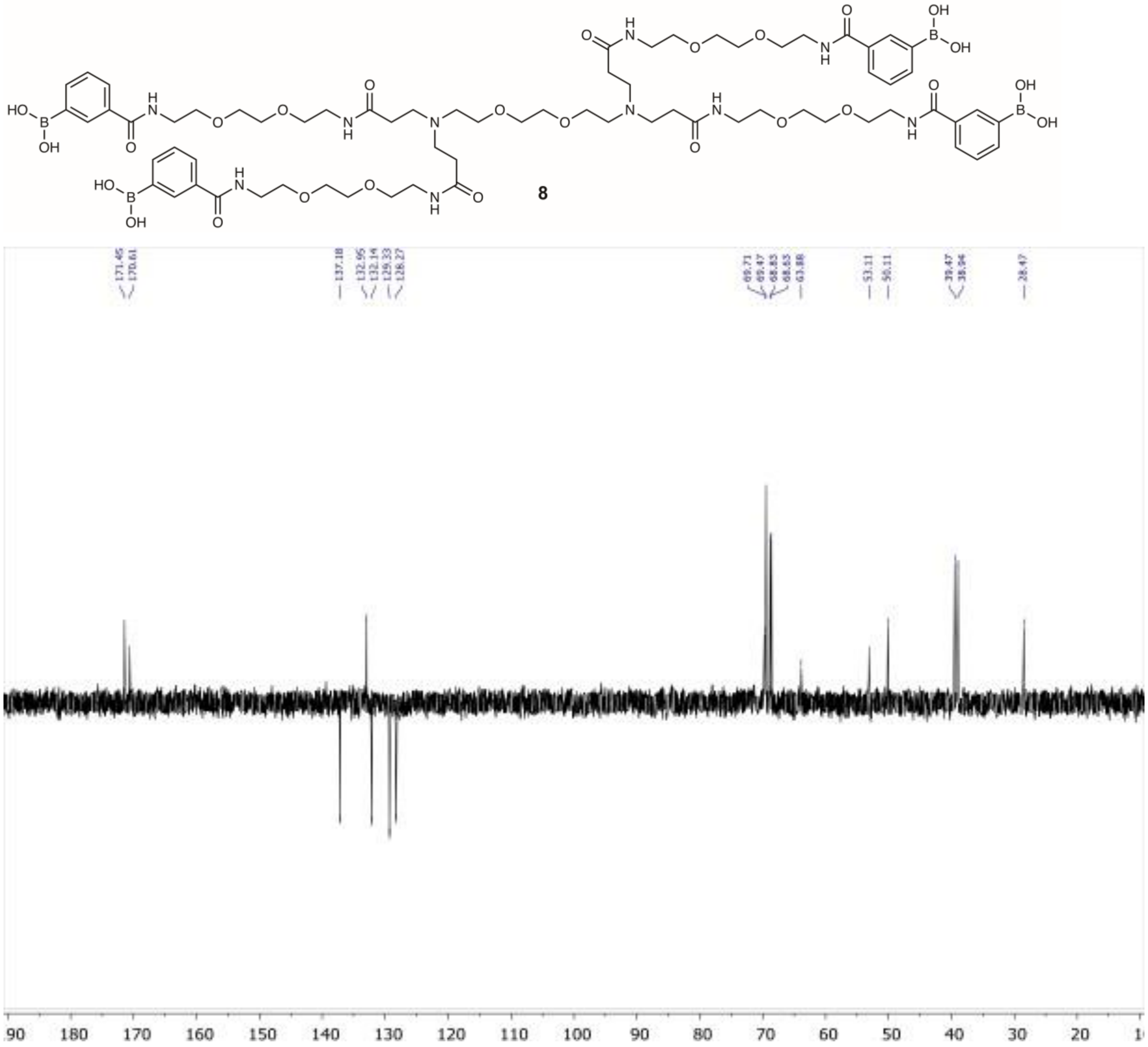


**Figure S20**. $^{13}$C DEPT NMR spectrum of compound **8**

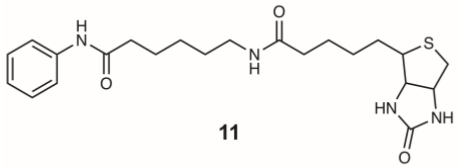

**11**

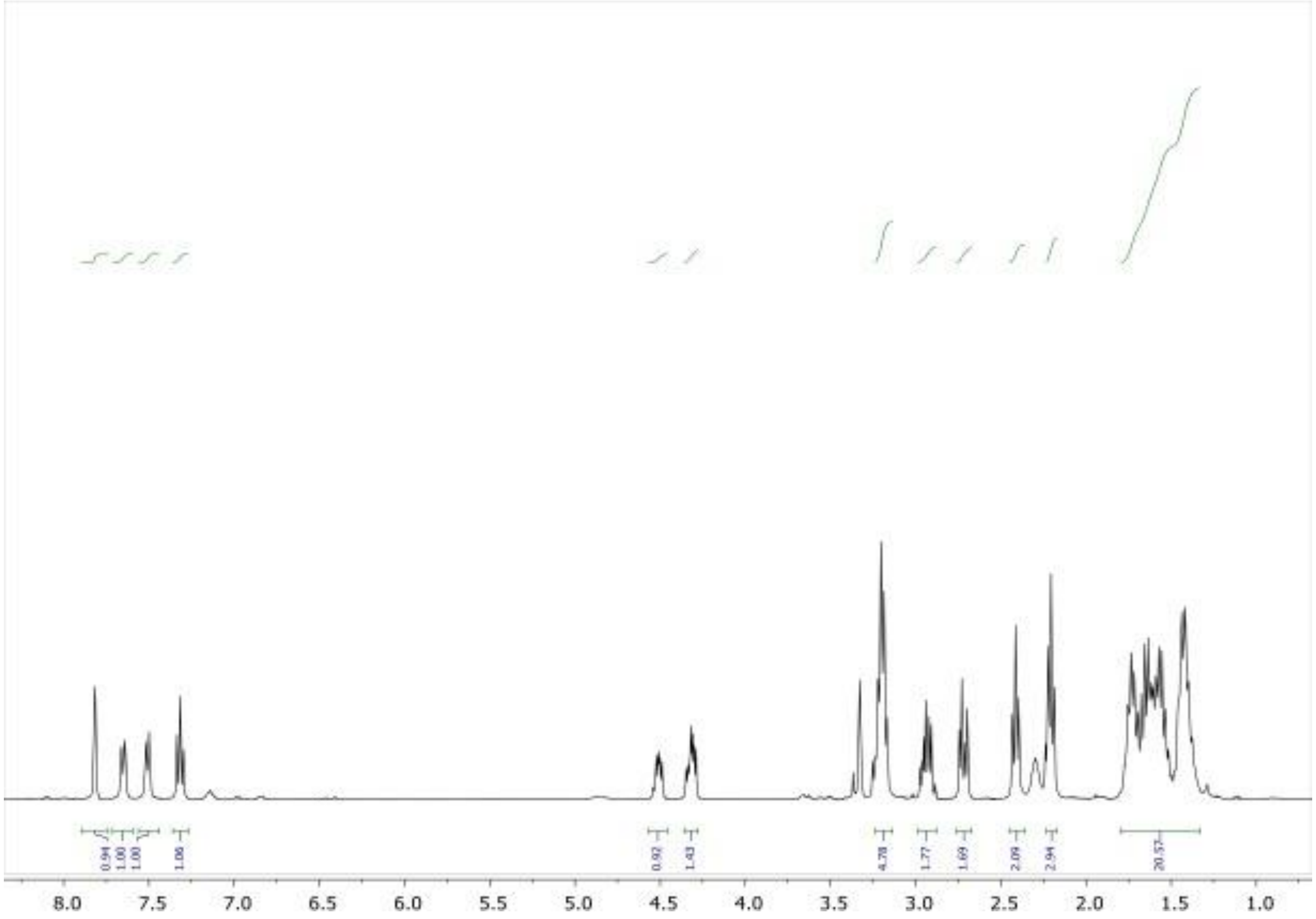

**Figure S21**. $^{1}$H NMR spectrum of compound **11**

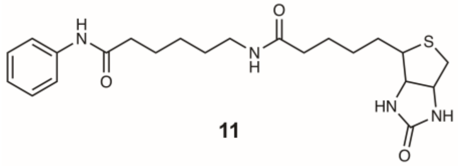


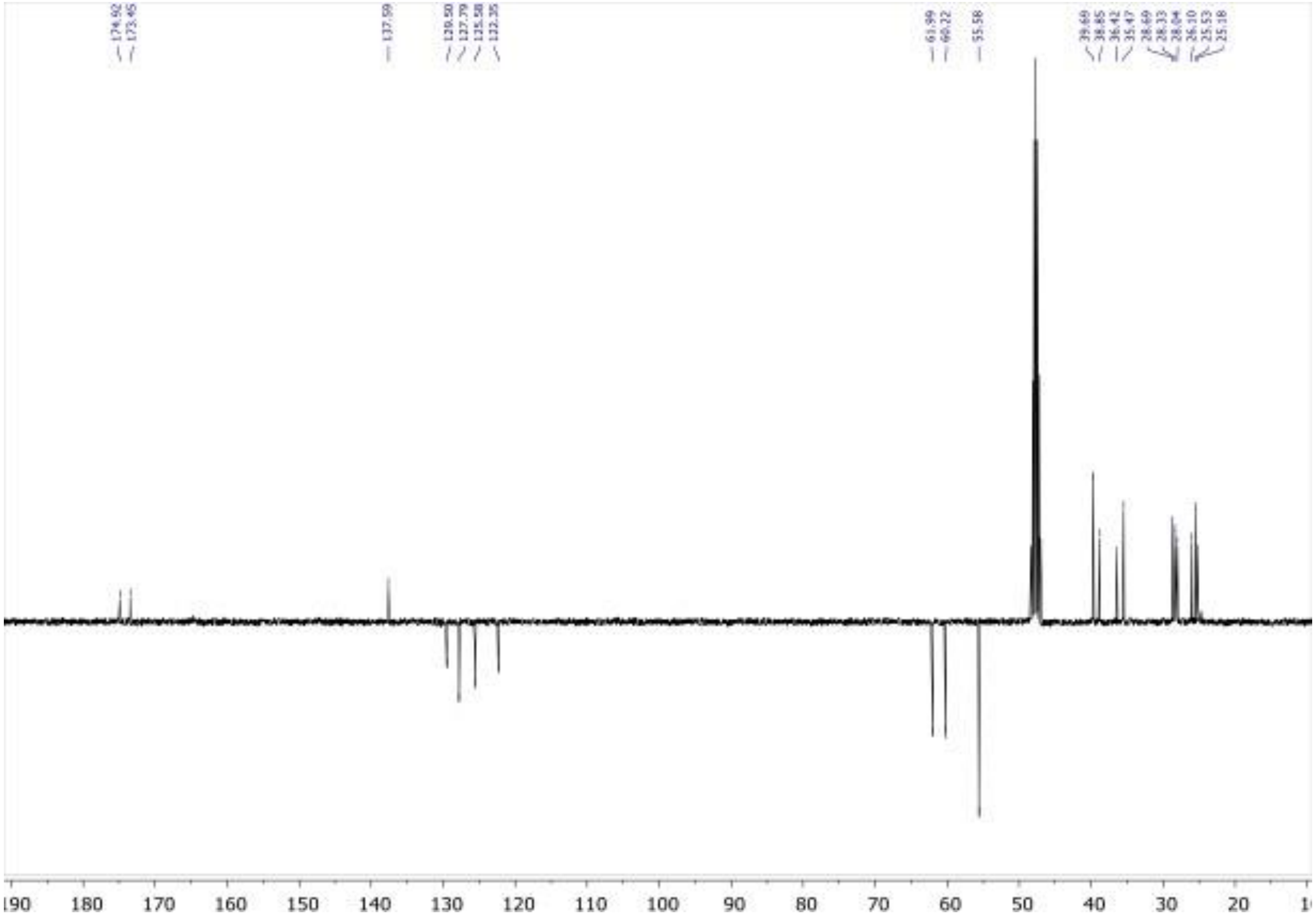


**Figure S22**. $^{13}$C DEPT NMR spectrum of compound **11**
